# Action prediction error: a value-free dopaminergic teaching signal that drives stable learning

**DOI:** 10.1101/2022.09.12.507572

**Authors:** Francesca Greenstreet, Hernando Martinez Vergara, Yvonne Johansson, Sthitapranjya Pati, Laura Schwarz, Stephen C Lenzi, Matthew Wisdom, Alina Gubanova, Fred Marbach, Lars Rollik, Jasvin Kaur, Theodore Moskovitz, Joseph Cohen, Emmett Thompson, Troy W Margrie, Claudia Clopath, Marcus Stephenson-Jones

## Abstract

Animals’ choice behavior is characterized by two main tendencies: taking actions that led to rewards and repeating past actions. Theory suggests these strategies may be reinforced by different types of dopaminergic teaching signals: reward prediction error (RPE) to reinforce value-based associations and movement-based action prediction errors to reinforce value-free repetitive associations. Here we use an auditory-discrimination task in mice to show that movement-related dopamine activity in the tail of the striatum encodes the hypothesized action prediction error signal. Causal manipulations reveal that this prediction error serves as a value-free teaching signal that supports learning by reinforcing repeated associations. Computational modeling and experiments demonstrate that action prediction errors alone cannot support reward-guided learning but when paired with the RPE circuitry they serve to consolidate stable sound-action associations in a value-free manner. Together we show that there are two types of dopaminergic prediction errors that work in tandem to support learning.

## Introduction

When animals and humans make repeated choices, they show two main tendencies: taking actions that led to rewards and repeating actions that they have taken in the past (Thorndike, 1911; Wood and Runger, 2016). The latter strategy is unrelated to value and could be seen as maladaptive, but recent work has shown there are computational advantages to learning perseverative behaviors (Gershman, 2020; Lai, 2021). Specifically, learning to repeat actions reduces the complexity of the policy, i.e., the mapping from states to actions. This reduction in policy complexity means it’s easier to store and remember what actions have been taken in the past as opposed to storing the value of actions in a particular context. Theoretical work has proposed that habitual associations are learnt through action repetition rather than through value-based learning (Dickinson, 1985; Miller et al., 2019; Wood and Runger, 2016). In line with this people default to habitual “simpler” strategies when their cognitive resources are taxed (Wood, 2014). Associations learned through action repetition have also been suggested to be more stable as they are not sensitive to changes in outcome contingency (Kim and Hikosaka, 2015). While the advantages of learning through repetition are beginning to be appreciated the mechanism for how these value-free associations form is unknown.

Dopamine neurons that encode reward prediction error (RPE), the difference between the reward obtained and the reward that was expected (Schultz et al., 1997), provide a critical teaching signal for value-based learning. These dopamine signals modify glutamatergic synaptic inputs to the striatum that were active prior to an unexpected reward (Gerfen and Surmeier, 2011; Reynolds et al., 2001; Reynolds and Wickens, 2002). In turn, these synaptic modifications increase the likelihood that actions that led to reward are repeated (Bamford et al., 2018; Costa, 2007; Cox and Witten, 2019). A recent model postulated that in value-free learning the strength of repetitive habitual associations should be updated by a movement-based and not a value-based teaching signal (Miller et al., 2019). The hypothesized teaching signal encoded an *action prediction error*, the difference between the action that is taken and the extent an action was predicted in a given state. Although the experimental evidence for this teaching signal is lacking, more theoretical work has shown how this value-free learning system could be implemented in the basal ganglia alongside the canonical value-based learning circuitry (Bogacz, 2020).

These models predict that there should be two types of dopaminergic teaching signals (Bogacz, 2020; Miller et al., 2019). A value-based RPE that teaches animals to repeat actions that lead to reward and a movement-based action prediction error that teaches animals to repeat the actions they have taken in the past. Interestingly, there are two main types of dopaminergic responses. Dopaminergic neurons located in the ventral tegmental area (VTA) and medial parts of the substantia nigra pars compacta (SNc) encode RPE (Cohen et al., 2012; Engelhard et al., 2019). On the other hand, dopaminergic neurons in the lateral SNc and substantia nigra pars lateralis (SNl) prominently respond to movement (da Silva et al., 2018; Dodson et al., 2016; Howe and Dombeck, 2016; Parker et al., 2016). The current prevailing theory is that movement-related dopamine release may trigger the initiation of action and modulate movement vigor (Klaus et al., 2019). In line with this, optogenetic stimulation of the SNc dopamine release can trigger the initiation of movement and increase movement vigor (da Silva et al., 2018; Howard et al., 2017; Howe and Dombeck, 2016). However, more recent experiments using optogenetic stimulation parameters that were calibrated to mimic physiological levels of SNc dopamine release did not trigger movement or lead to an invigoration of ongoing action (Coddington and Dudman, 2018). In contrast, these stimulation parameters were sufficient to act as a teaching signal to drive learning. Although none of these experiments specifically manipulated movement-related dopamine release, the recent experiments are compatible with the idea that movement-related dopamine release could act as an alternative teaching signal rather than as a driver of action initiation (Coddington and Dudman, 2019).

Our aim was to determine if movement-related dopaminergic activity encodes an action prediction error and serves as a value-free teaching signal to reinforce repeated state-actions associations. To address this, we examine the role dopamine plays in the tail of the striatum (TS) as mice learn an auditory discrimination task called the “cloud-of-tones” task (Znamenskiy and Zador, 2013). In this task, selective corticostriatal plasticity develops between the auditory cortex and the TS in both rats and mice (Ghosh and Zador, 2020; Xiong et al., 2015). These changes are one of the best examples of experience-dependent corticostriatal modifications, but they occur in a region of the striatum that does not receive dopamine RPE signaling (Watabe-Uchida and Uchida, 2018). Rather, the dopaminergic activity in this region, arising from neurons in the SNl, is modulated by movement and other non-reward related processes (Dodson et al., 2016; Menegas et al., 2018; Menegas et al., 2017). Here we show that dopaminergic input to this region is important for learning, and that dopaminergic activity during the task is movement-related. We go on to show that this movement-related dopaminergic activity encodes an action prediction error. Causal manipulations show that this value-free teaching signal can reinforce state-action associations, essentially biasing mice to repeat the actions that they have taken in the past. We propose a model where this movement-based teaching signal works in conjunction with canonical RPE signaling to boost and stabilize learning about consistent state-action associations.

## Results

### TS is needed for efficient learning and expression of the auditory discrimination task

We trained mice on the “cloud-of-tones” task (Znamenskiy and Zador, 2013). Mice initiate trials in the center port, where a low or high frequency auditory stimulus indicates which port (left or right) will deliver a water reward (Figure 1A). To determine if the TS was required to perform this frequency-discrimination task, we bilaterally inactivated the TS with injections of muscimol (GABA-A receptor agonist) in expert mice. In line with previous reports (Guo et al., 2018), inactivation of the TS impaired task performance (Figures 1B and 1C). In contrast, inactivation of the dorsomedial striatum (DMS), which also receives projections from primary auditory areas (Hunnicutt et al., 2016), had no significant impact on performance. We next set out to determine if both types of projection neurons in this region, known as dopamine D1 and D2 receptor-expressing striatal projection neurons (SPNs), were needed to perform this task. To examine their respective contributions to performance we used optogenetics to unilaterally inactivate either D1 or D2 SPNs in trained mice (Figure 1D and Figures S1A and S1B). Optogenetic inactivation of either population had a significant effect on task performance. Inactivation of D1 SPNs resulted in an ipsilateral choice bias relative to the stimulated hemisphere, while inactivation of D2 SPNs resulted in a contralateral choice bias (Figure 1E and 1F). Together, these results show that in agreement with recent results the TS is needed to execute the learned behavior and that both populations of SPNs in the TS exert opposing contributions to the auditory-guided choices.

**Figure 1:**
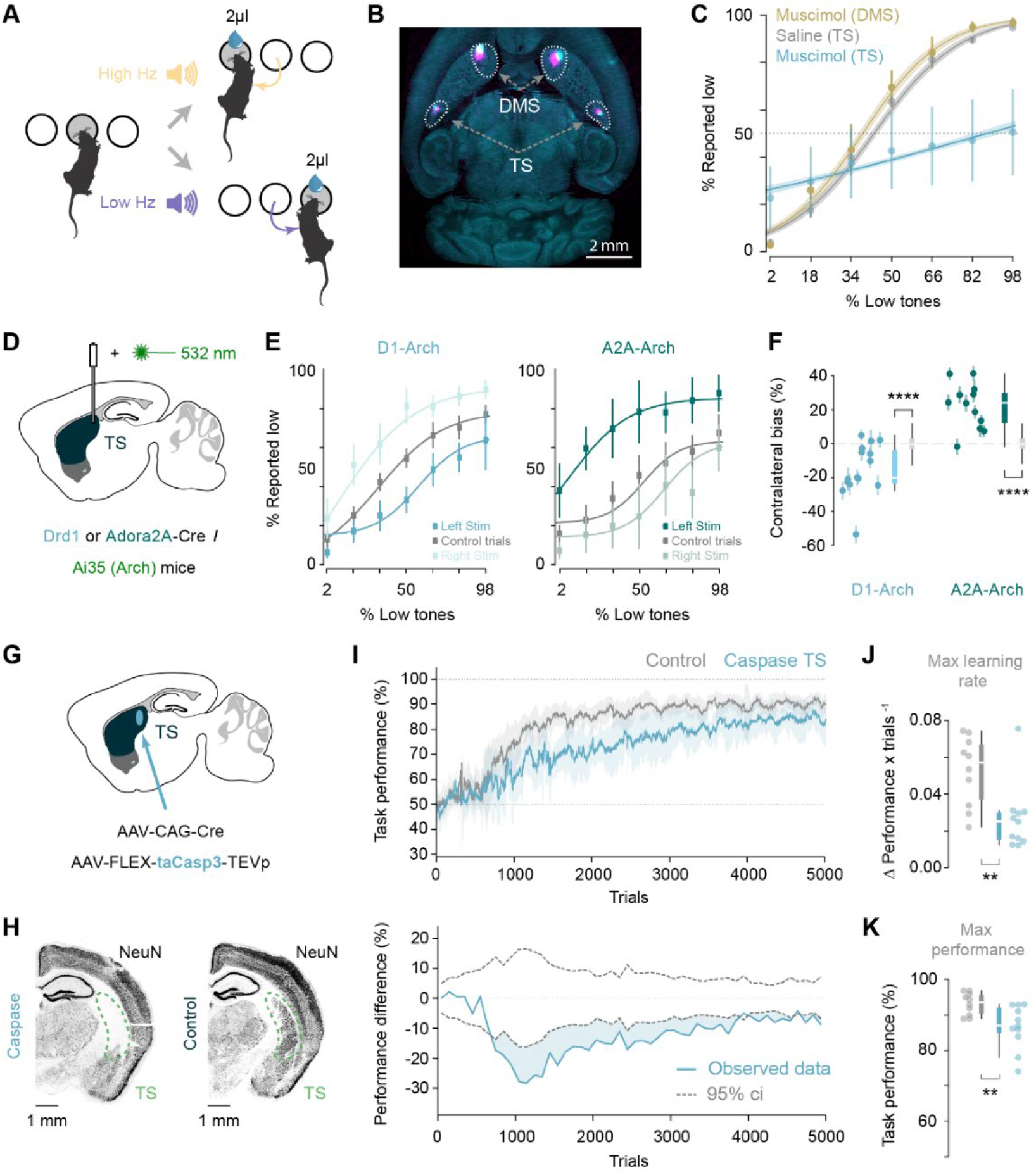
TS is needed for learning and expression of the auditory discrimination task. **A** - Schematic of the task. The mouse initiates trials in the center port. Low or high frequency stimuli predict water reward on either the right or the left port. **B** - Transverse section of a mouse brain showing the location of the muscimol injections in the DMS and TS. Bilateral muscimol injection locations (for 4 sessions in total) are indicated by the co-injection of fluorescent CTB (cyan and magenta for both the TS and DMS). **C** - Psychometric task performance of mice injected with saline in TS (2 mice; 4 sessions), and muscimol in DMS (3 mice; 11 sessions) and TS (5 mice; 15 sessions). Error bars represent the standard deviation across sessions, and the shaded area the standard deviation of the model fit. **D** - Schematic of the experimental approach for acute inhibition of D1 SPNs or D2 SPNs in the TS. **E** - Psychometric task performance of unstimulated and unilaterally (left or right hemisphere) opto-stimulated trials for the D1 Arch (8 mice; 16 sessions) and A2A Arch (8 mice; 12 sessions) mice. Error bars represent the standard deviation across sessions. **F** - Quantification of the bias (see methods) for each session shown in E. Scatter dots represent the means and standard deviation for each session, and coloured box plots represent the distribution of the means compared to the baseline in gray (see methods). D1 Arch: p=2.38×10^−5^; A2A-Arch: p=1.55×10^−7^; Kruskal-Wallis tests. **G** - Schematic of the experimental approach for lesioning the TS. **H** - Histology showing the neurons in TS for an example lesion and control mice. **I** - Top graph: learning rate (performance vs. trial number) of the lesioned TS and the control groups (shaded area indicates standard deviation). Bottom graph: differences in performance between the groups. Dotted lines indicate the 95% confidence interval for the shuffled data (see methods). **J** and **K** - Differences on the maximum learning rate (p=0.009, Kruskal-Wallis test) and maximum performance (p=0.006, Kruskal-Wallis test) between the groups.

To test if the TS was also needed to learn the task, we ablated the TS prior to training using a viral-mediated Caspase-based strategy (Figures 1G and 1H, and Figures S1C and S1D). Lesions of the TS caused a deficit in learning (Figure 1I), reducing both the learning rate and the maximum performance reached on the task (Figure 1J-1K). Together, these results show that the TS is needed for learning and performing the auditory frequency-discrimination task.

### TS dopamine release is needed to efficiently learn the discrimination task

Next, we tested whether dopaminergic innervation of the TS was important for learning. We ablated TS-projecting dopamine neurons using a selective neurotoxin, 6-hydroxydopamine (6-OHDA). Injections of 6-OHDA into the TS strongly reduced dopamine innervation of this region but not the adjacent dorsal striatum (Figure 2A and S2A). Recapitulating the general TS lesions effects (Figure 1I), TS-dopamine ablated mice had a deficit in learning (Figure 2B). These mice had reduced learning rates (Figure 2C) as well as a lower maximum performance reached on the task (Figure 2D). Across individual mice, the deficit in learning was significantly correlated with the size of the dopaminergic lesion (Figure 2E). In contrast to the learning deficits, 6-OHDA lesions affected neither the time taken to move from the center port to the choice ports (Figure S2B) nor the time taken between trials (Figure S2C). These results demonstrate that dopaminergic innervation of the TS facilitates learning the frequency-discrimination task but does not affect movement initiation or vigor.

**Figure 2:**
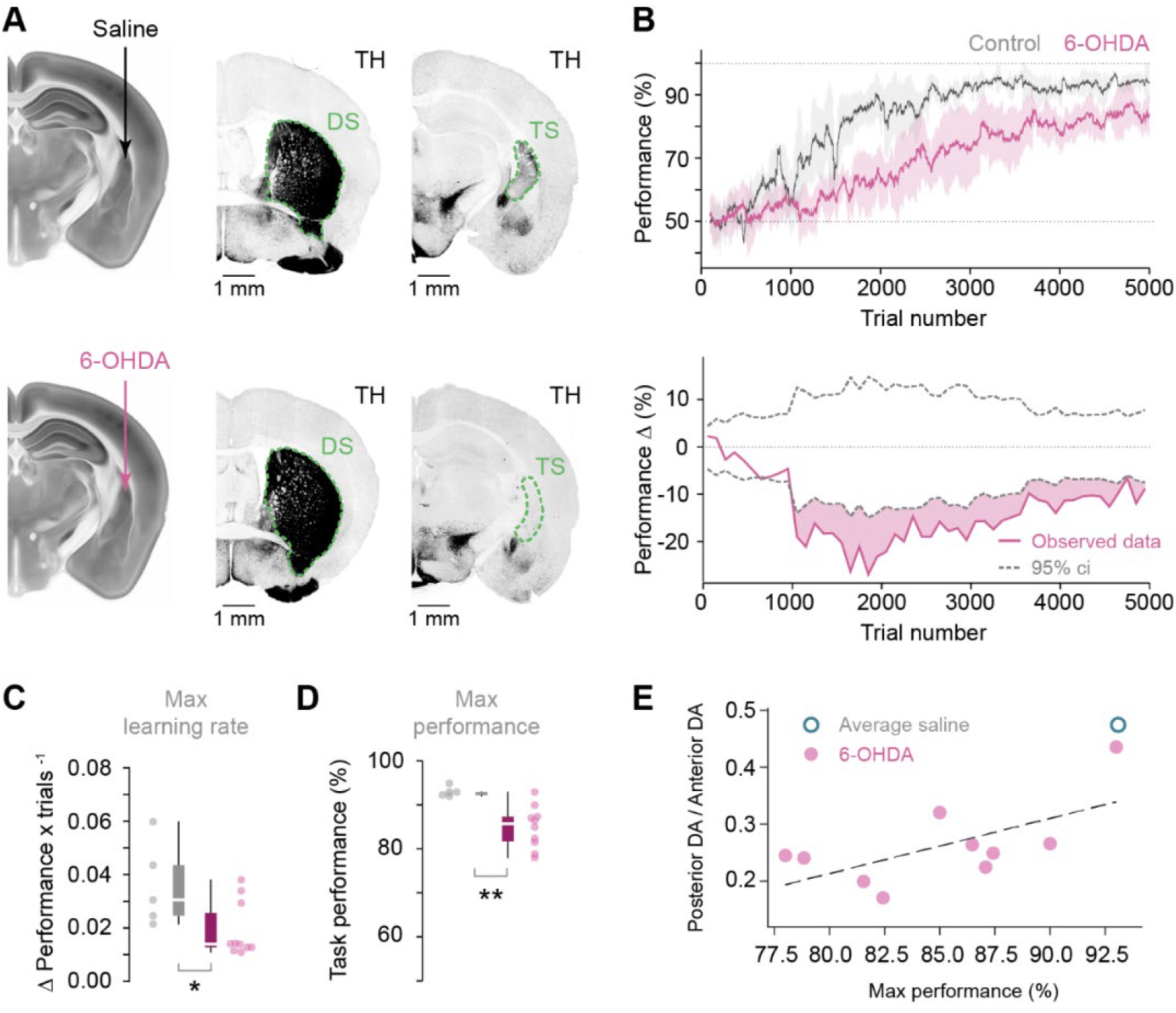
TS dopamine release is needed for efficient learning of sound discrimination. **A** - Indication of injection sites in coronal slices of the reference atlas, and example TH-stained (dopamine axons) coronal slices at the level of the dorsal striatum (DS) and the TS for control and lesion mice. **B** - Top graph: learning rate (performance vs. trial number) of the TS dopamine-ablated and the control groups. Bottom graph: differences in performance between the groups. Dotted lines indicate the 95% confidence interval for the shuffled data (see methods). **C** and **D** - Differences on the maximum learning rate (p=0.019, Kruskal-Wallis test) and maximum performance (p=0.002, Kruskal-Wallis test) between the groups. Error bars indicate standard deviation. **E** - Correlation between the maximum performance achieved for each mouse and the lesion size (p=0.049, two-sided Wald test).

### Dopamine in TS is correlated with contralateral movement

Dopamine activity in the SNl-TS pathway is activated by movement and other non-reward related processes (Dodson et al., 2016; Menegas et al., 2018; Menegas et al., 2017). To understand the role TS dopamine plays in the auditory discrimination task, we measured its dynamics with dLight1.1, a genetically encoded fluorescent dopamine sensor (Patriarchi et al., 2018) (Figure S3). Dopamine signals were monitored during the initial learning phase with fiber photometry and compared to the canonical reward prediction error signals in the ventral striatum (VS) (Figure 3A-C). We observed that the biggest TS dopamine responses correlated in time with movements from the center port to the port contralateral to the recording fiber, in sharp contrast to the large reward responses in VS (Figure 3D-E).

**Figure 3:**
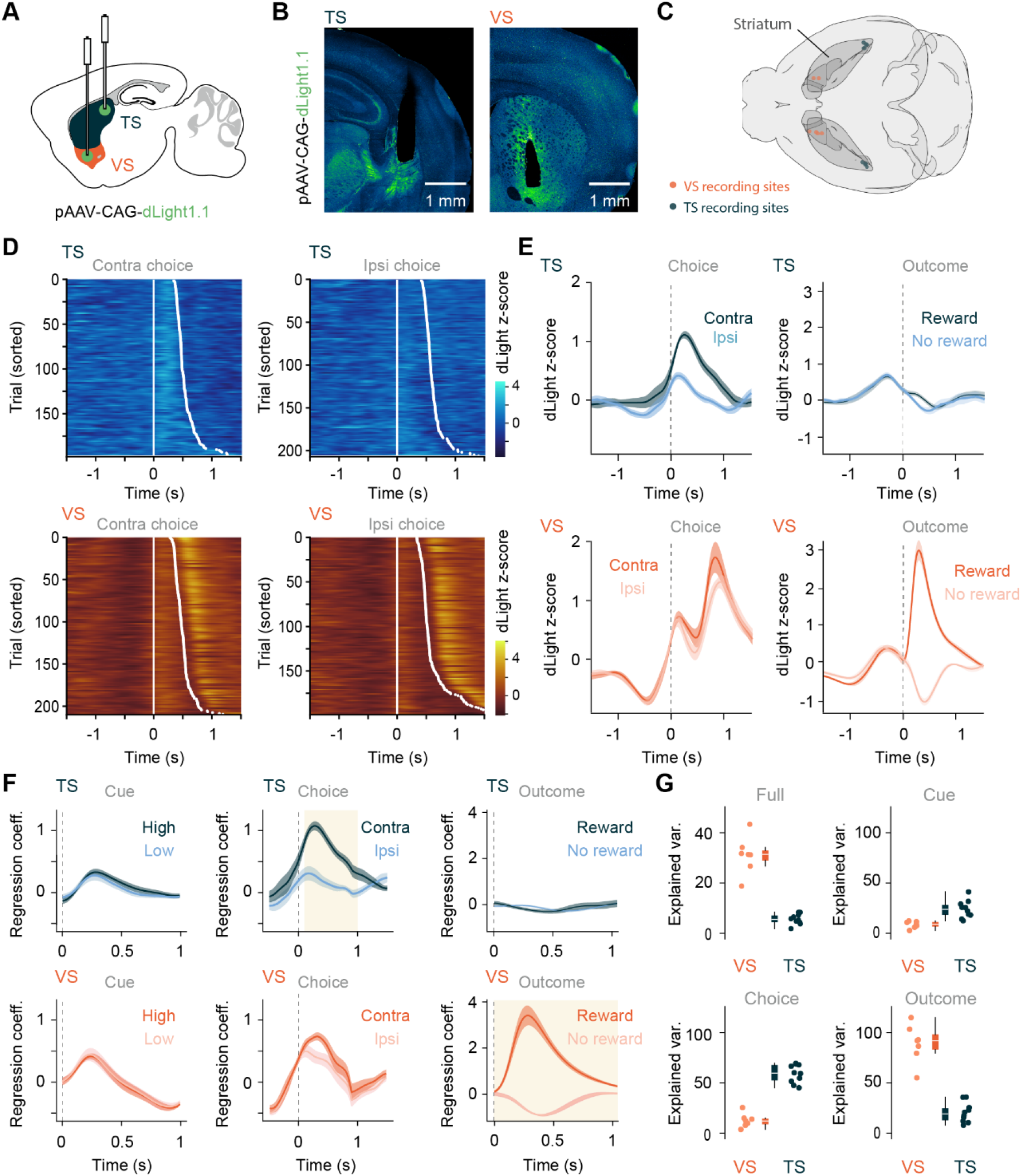
Dopamine in TS is correlated with contralateral movement. **A -** Schematic of the experimental approach for recording dopamine activity in the VS and TS. **B -** Fluorescent images showing optical fiber locations and dLight expression (green) in the TS (left) and the VS (right). **C -** Top down view of 3D brain render showing the recording fiber locations in the VS and TS. **D -** Example of a TS (blue) and VS (orange) recording session aligned to time of leaving the center port to make a contralateral choice (left column) or an ipsilateral choice (right column). White dot shows time of entering contralateral or ipsilateral choice port. **E -** Average photometry traces in TS (n=9 mice) and VS (n=7 mice) aligned to choice, time leaving the center port, for contralateral and ipsilateral choices (left column) and aligned to reward outcome, time entering choice port, (right column). Lines represent the average response profile across mice. **F -** Average response kernels to behavioral events for recordings in the TS (blue) and VS (orange). Lines represent the mean kernels across mice. Shaded time windows show significant differences between the two kernel types in each subplot, calculated by performing two-sample t-tests on 0.1 s bins and a p-value threshold for significance of 0.01. **G -** Percentage explained variance of the whole recording session (VS median = 31.3; TS median = 5.50) and for different behavioral kernels for linear regression models fitted on VS and TS recordings. All error bars represent SEM.

To better separate the dopamine response associated with individual behavioral events, which may have overlapping temporal dynamics, we fitted a linear regression model to photometry data obtained early in training. The regression model included three types of behavioral events: cue (center port entry), choice (center port exit) and outcome (side port entry). The model produced a predicted response kernel for each of these types of behavioral events which captured the stereotypical modulations of the dopamine response for each behavioral event (Figure 3F). The response in the VS was best explained by the outcome kernel, which captured the large dopamine responses observed when reward was available (Figure 3E-G). This response was significantly different to unrewarded outcomes, characterized by a dip in VS dopamine levels (Figure 3E and 3F). In contrast, there was very little dopaminergic response in the TS to either reward or non-rewarded trials, with no significant difference between these conditions (Figure 3E-G). Instead, in the TS, the kernel with the largest dopaminergic response was the movement-locked activity (Figure 3E-G), with contralateral movements (the opposite direction to the recorded hemisphere) eliciting a significantly larger response than ipsilateral movements (Figure 3F). The movement-locked activity in the VS was smaller, with no significant difference between contra and ipsilateral actions (Figure 3F and 3G). In both the TS and VS there was a small response at the time of the cue that did not differ between high or low frequency sound trials. Together, these results show that VS dopamine activity significantly encodes reward outcome, consistent with reward prediction error, while the TS dopamine activity encodes information about movement.

To confirm that the TS dopamine activity is movement and not sensory related, we omitted the sound cue on a subset of trials. On these trials mice still moved to one of the two choice ports but there was no sound to guide their action (Figure S4A). On the silent trials there was still a comparable TS dopamine response to contralateral movements that was not significantly different from the control trials where the sound was on (Figure S4B-D). This result shows that the movement-related dopamine response is not driven by or dependent on the auditory cue. To determine if the movement related activity is task dependent, we recorded TS dopamine activity as mice freely explored an open arena. As with our recordings in the task, TS dopaminergic activity rose just prior to contralateral orienting movements and peaked following movement initiation (Figure S4E-J). Finally, if the task-related dopamine activity in the TS is driven by movement then the size of the signal should correlate with degree of movement. To determine if that was the case, we used behavioral tracking to measure the cumulative turn angle for every trial and then grouped trials into quartiles (Figure S4K-N). There was a significant correlation between the turn angle and the TS dopamine activity, with larger turn angles resulting in larger TS dopamine responses on average (Figure S4O and S4P). This correlation between degree of movement and dopamine activity was not present in the VS (Figure S4P). Together these results confirm that the TS dopaminergic activity encodes information about movement and not sensory or reward related information.

### TS dopamine release decreases in magnitude during learning

Taken together, the results so far show that dopamine innervation of the TS is important for learning the auditory frequency-discrimination task and that dopaminergic activity in the TS encodes information about movement and not reward. While there is rich theoretical work about how dopamine RPEs can drive corticostriatal plasticity and learning (Cox and Witten, 2019; Klaus et al., 2019; Reynolds et al., 2001; Schultz et al., 1997) how movement-related dopamine contributes to learning is unknown. One possibility, inspired by recent computational models of habit formation, is that movement-related dopamine may encode a value-free prediction error that works in concert with RPEs to support stable “habitual” learning (Bogacz, 2020; Miller et al., 2019). In these models there are two ways for the striatum to control behavior. Firstly, a value-based system that learns to predict the reward associated with actions in a particular state. These reward predictions are learnt and updated by dopaminergic RPEs. Secondly, a value-free system that learns to predict the most likely action to be taken in a particular state. These value-free action predictions are updated by an action prediction error (APE), which encodes the difference between the action that is taken and the extent to which an action was predicted in a given state.

If the movement-related dopaminergic activity in the TS encodes an APE, then the first prediction is that movement-related dopamine activity in the TS should initially be large and then decrease over time as mice learn the task. As early in learning there is no action predicted by the auditory cues and therefore no subtraction from the action that is taken, this should result in a large movement-related dopamine signal. However, as learning occurs the mice start to predict the action they will take in response to a particular stimulus, leading to a decrease in APE over time. To formalize and test this prediction we recorded TS and VS dopamine signals during the full extent of mice learning the task and compared these signals to values of RPE and APE from a dual value-based/value-free reinforcement learning model (Figure S5A-C). The task was modeled as a set of discrete states. In the value-based system the value of these states is updated using RPEs (Figures S5A and S5B). The strength of the state-action associations in the value-free portion are updated by APEs (Figure S5A). In line with encoding an APE signal, the movement-related dopamine activity in the TS decreased as mice learned the task (Figure 4A-D). The dopamine response was largest in naive mice and then decayed over thousands of trials as mice became experts at the task, matching the pattern of exponential decay of the value-free system of the model (Figure 4C). In contrast and in line with the value-based system, dopamine recordings in the VS showed that the cue-related dopamine activity grew over time (Figure 4E-H), characteristic of a RPE signal. These results show that the movement-related dopamine activity in the TS does not reflect a pure movement signal that is invariably linked to action initiation, rather it decays as mice come to predict the action that will be taken in a certain state, as expected from an APE signal.

**Figure 4:**
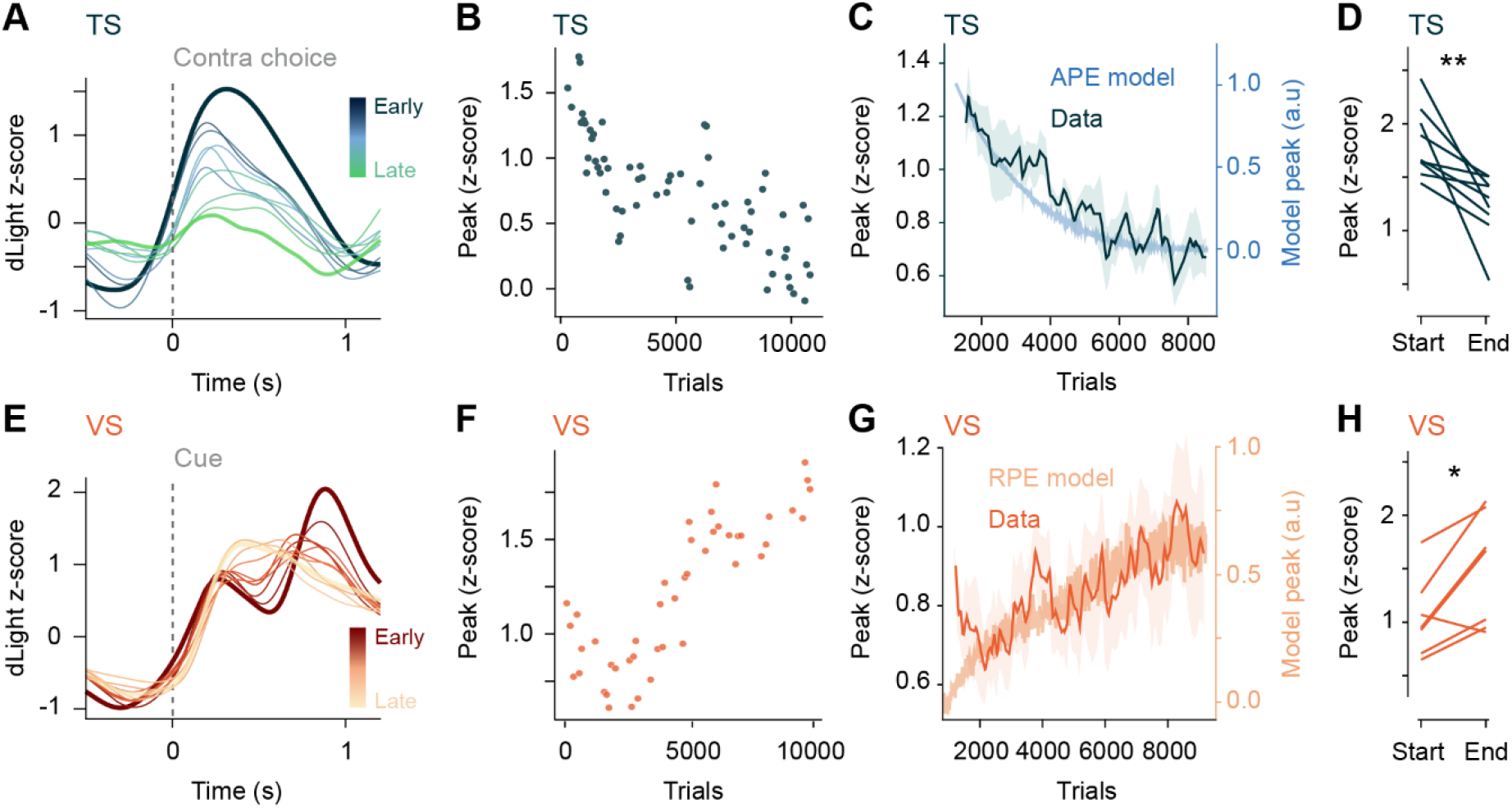
TS dopamine release decreases in magnitude during learning. **A -** Example dLight recording over time in the TS aligned to contralateral choice. Each trace is the average of 200 trials, with each trace color-coded by the stage in learning it was recorded from early-late. The traces from the earliest (first recorded session) and latest (last recorded session) traces are represented with a thicker line. **B -** Example TS dopamine response size to contralateral choice over training binned every 50 trials. **C -** Average change in contralateral choice-aligned TS dopamine response across training. The solid dark line represents the mean across mice (n=6 mice), shaded error bars represent SEM. Light blue trace is the peak action prediction error response to contralateral choice over training from an average of 100 model agents. **D -** Comparing size of contralateral choice-aligned dopamine response in TS in the first and last session of training (n=9 mice), p=0.006 (two related samples t-test). **E -** same as A but for VS recordings and aligned to the contralateral cue. **F -** Same as B but for VS recordings aligned to the contralateral cue. **G -** Same as C but for VS recordings (n=7 mice). Light orange trace is the peak reward prediction error response to contralateral cue over training from an average of 100 model agents. **H -** Same as D but for VS recordings (n=7 mice), p=0.016 (two related samples t-test).

### TS dopamine release is state-dependent and value-free

The learning-associated decrease of the movement-related TS dopamine signal suggests that it is modulated by how predictably an action is taken in response to a particular state (auditory cue). Therefore, if trained animals were to make a familiar movement in response to an unfamiliar sound the action would not be predicted by the stimulus and consequently the resulting APE should be larger. To examine this hypothesis, we tested expert mice on a variation of the task, in which we replaced the auditory cue associated with the contralateral action with a novel cue (white noise) (Figure 5A and 5B). The presentation of the new stimulus decreased the probability of mice choosing the appropriate port (Figure S5D) and led to a significant increase of the movement-related TS dopamine signal (Figure 5C and 5D). This is expected from an APE signal, as the action is no longer predicted by the stimulus and therefore not subtracted from the action signal (Figure 5B). In contrast, and predicted by the RPE model, the VS dopamine cue response to the novel sound was smaller (Figure 5E and 5F), as there is less value predicted by the novel cue (Figure 5B).

**Figure 5:**
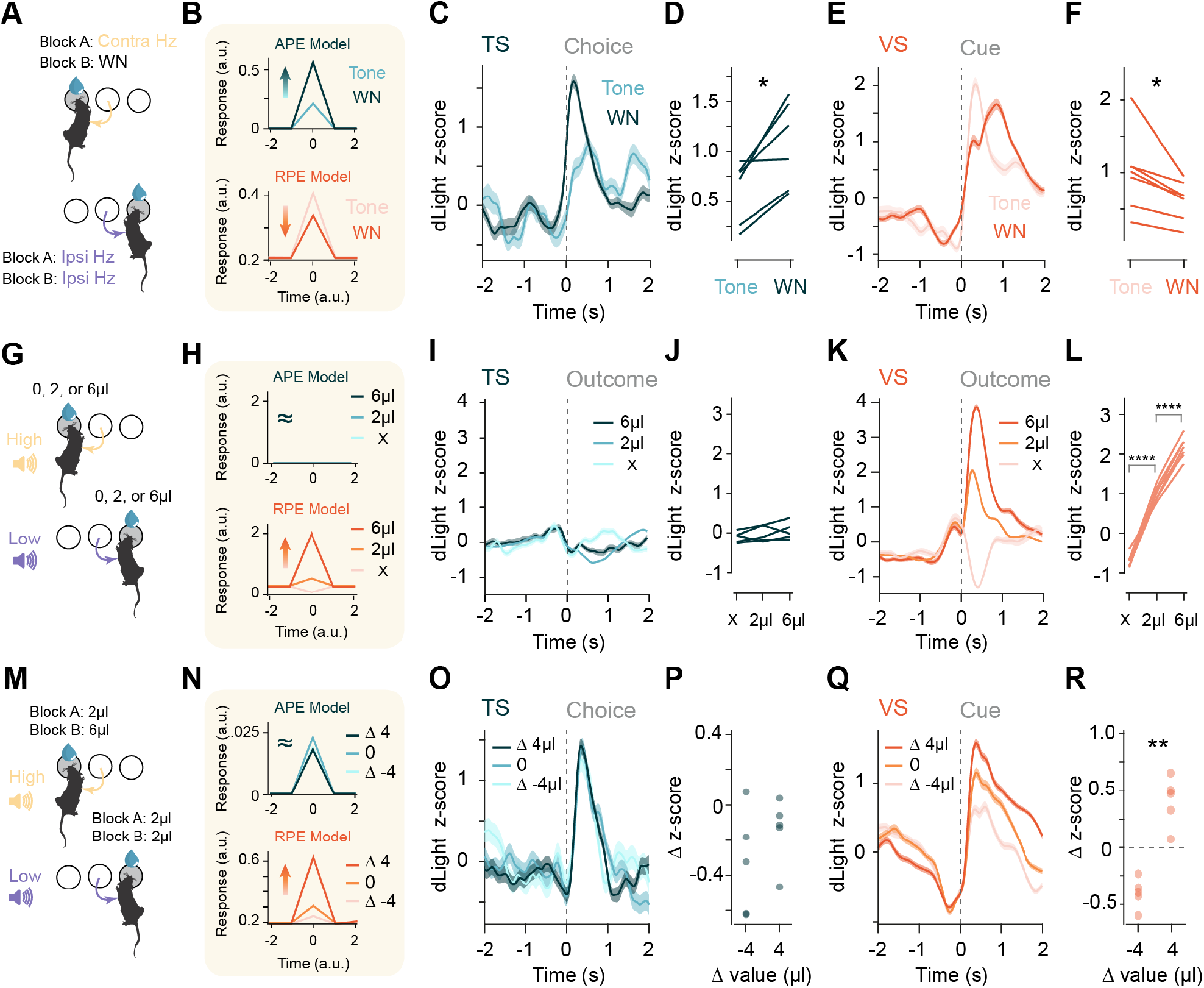
TS dopamine release is state-dependent and value-free. **A -** Schematic of the state change task design. In each case midway through the session the auditory cue associated with the action contralateral to the recording hemisphere was replaced with a novel white noise stimulus. **B -** Modeled responses for how APE and RPE signals would change following the state change. **C -** Example TS dopamine responses to the contralateral choice in response to the normal or the white noise cue. Solid line represents mean responses for one mouse across trials for each trial type. **D -** Group data comparing the TS dopamine response to the contralateral action before and after the introduction of the novel state (p=0.01, two related samples t-test) (n=6 mice). **E -** Same as **C** but for a VS recording aligned to cue. **F -** Same **D** but for VS recording aligned to cue. (p=0.02, Wilcoxon signed-rank test) (n=6 mice). **G -** Schematic of the outcome manipulation task design. **H -** Modeled responses for how APE and RPE signals would respond to changes in reward outcome. **I -** Example TS response to omissions, normal sized and large rewards. Solid line represents mean responses for one mouse across trials for each trial type. **J -** Group data (n=6 mice) for TS dopamine response size to omission, normal and large reward (p=0.84, p=0.54, two related samples t-test, adjusted using Bonferroni correction). **K -** Same as **I** but for a VS recording. **L -** Same as **J** but for VS recording (n=7 mice, p=3.16×10^−6^, p=8.87×10^−6^, two related samples t-test, adjusted using Bonferroni correction). **M -** Schematic of the predicted value manipulation task design. For each mouse the value of the left and right port was altered in separate sessions. **N -** Modeled responses for how APE and RPE signals would respond to changes in predicted outcome value. **O -** Example movement-aligned TS response when the relative value of the cues changed. **P -** Summary data showing the change in movement-aligned response in the TS when the relative value of the contralateral choice decreases (- 4) or if the relative value increases (+4) (p=0.26, two related samples t-test) (n=5 mice). **Q -** Same as **O** but for the VS response aligned to cue. **R -** Same as **P** but for the VS response aligned to cue (p=2.41×10^−4^, two related samples t-test) (n=5 mice). Data are represented as mean ± SEM in C, E, I, K, O, Q.

Another hallmark of an APE is that it should be value-free and should not be modulated by either the outcome or the predicted value of an action. To test this, we first varied the reward outcome, either delivering a larger than expected reward or omitting reward on a subset of trials (Figure 5G). In line with encoding an APE, the dopamine signal in the TS did not significantly respond to a larger or smaller than predicted reward (Figure 5H-J). In contrast, the VS dopamine signal was altered in a manner consistent with RPE (Figure 5H, 5K and 5L). Next, to test if the TS dopamine signal is modulated by changes in the predicted value of an action, we increased the reward associated with one port in a block-wise fashion (Figure 5M). This caused a small but significant behavioral bias in the choice of the mice towards the high value port (Figure S5E), but it did not influence the size of the TS dopamine signal (Figure 5N-P). In contrast to the signal in the TS, the predicted value manipulation altered the VS dopamine signal in a manner consistent with RPE (Figure 5N, 5Q and 5R).

Finally, to determine if the modulation in the movement-related TS dopamine signal was consistent with other theoretical models of dopamine activity, we modeled how the signal would appear if it encoded movement *per se*, novelty, or salience (see methods). Unlike the models for APE and RPE, none of the models consistently matched the pattern of modulation across conditions that we observed in the dopamine activity recorded in the TS or VS (Figure S5). Taken together, our results reveal that the movement-related TS dopamine signal is consistent with encoding an action prediction error.

### TS dopamine release reinforces state-action and not state-outcome associations

In computational models of habit formation, the APE signal is used as a value-free teaching signal to update the strength of state-action associations (Bogacz, 2020; Miller et al., 2019). To determine if the TS dopamine signal also serves this function, we optogenetically stimulated TS dopamine release at different epochs of the frequency-discrimination task (Figure S6A and S6B), using stimulation parameters that have been shown to mimic endogenous dopamine release (Coddington and Dudman, 2018). First, to mimic the endogenous movement-related TS dopamine signal, we stimulated unilaterally, at the center choice port, in trials where the cue signaled a contralateral choice (Figure 6A and 6B). In the block in which this optogenetic stimulation was active, animals displayed a bias towards contralateral choices, as would be expected if the contralateral state-action association was strengthened (Figure 6C and 6D). Optogenetic stimulation did not bias action directly as there was no choice bias on individually-stimulated trials (Figure S6C-E). Stimulating dopamine release in the VS at the time of choice had no effect on the animal’s choice bias (Figure 6D). Second, to determine if TS dopamine release at the time of choice outcome also had an effect, as has previously been reported (Menegas et al., 2018), we bilaterally stimulated TS dopamine release whenever mice poked into the selected side port (Figure 6E). Unlike stimulation at the time of choice, stimulation at the time of outcome did not bias the mice to repeat or avoid the choice that led to the optogenetic stimulation (Figure 6F). These results show that TS dopamine release can, as predicted by the computational models, reinforce state-action but not state-outcome associations.

**Figure 6:**
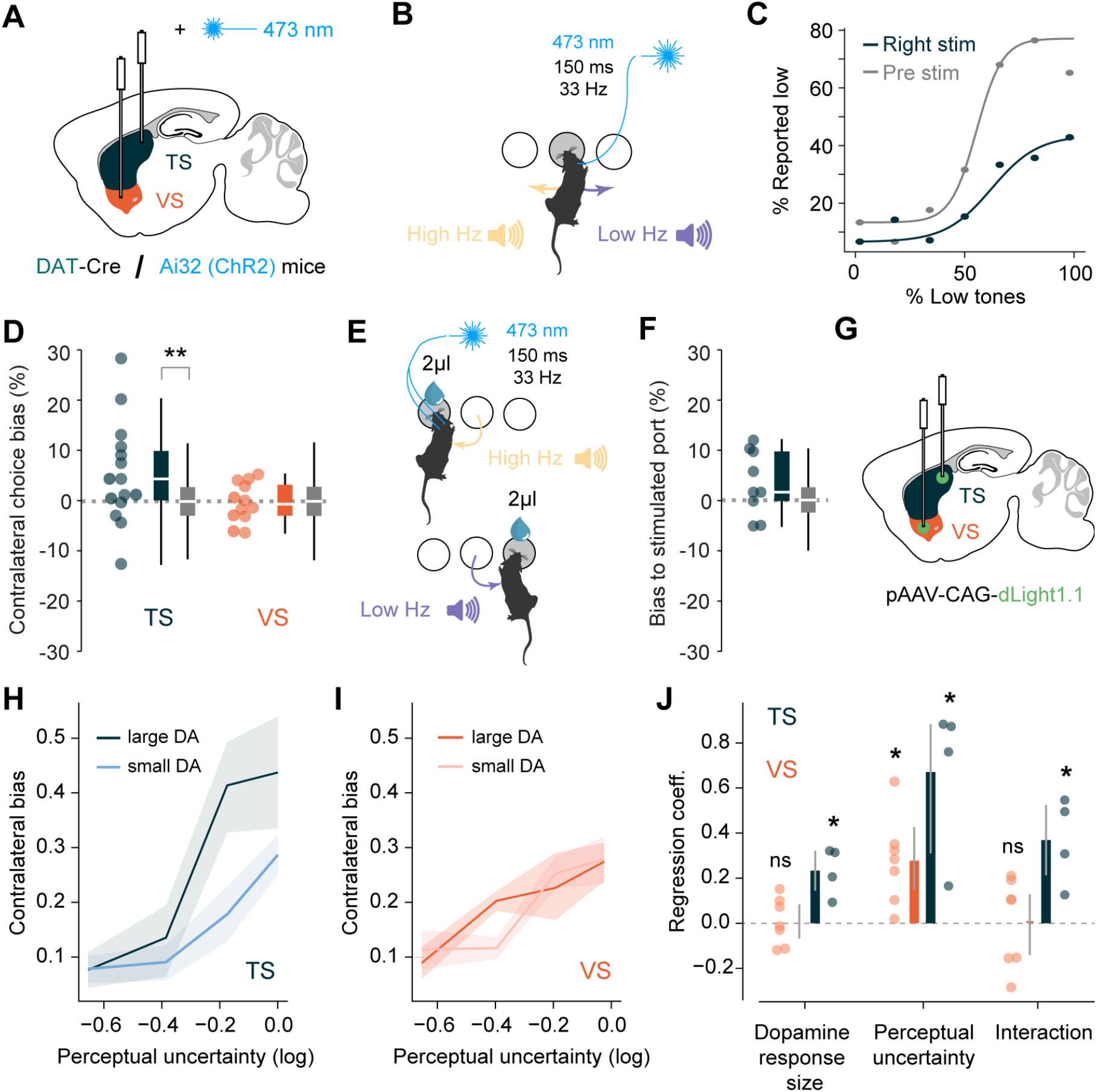
TS dopamine release reinforces state-action and not state-outcome associations. **A** - Schematic of the experimental approach for stimulating dopamine axons in the VS and the TS. **B** - Schematic of the stimulation protocol for the state-action experiment. Mice were stimulated in the center port unilaterally for the contralateral-movement state. **C** - Example session for the stimulation of the TS. **D** - Distribution of session biases (both in scatter and colored box plots) for the TS and the VS stimulations of the state-action experiment, compared to the shuffled data (see methods) (TS: p=0.005; VS: p=0.57; Wilcoxon rank-sum test). **E** - Schematic of the stimulation protocol for the state-outcome experiment. Mice were stimulated in one of the side ports bilaterally. **F** - Same as D for the state-outcome TS-stimulation experiment (p=0.053; Wilcoxon rank-sum test). **G** - Schematic of the experimental approach for recording dopamine activity in the VS and TS. **H** - Average change across mice in the contralateral bias for trials preceded by small or large dopamine choice movement responses in TS. **I** - Average change across mice in the contralateral bias for trials preceded by small or large dopamine reward responses in the VS. **J** - Regression coefficients for linear regression model predicting contralateral choice bias from previous trial dopamine response size, current trial perceptual uncertainty and the interaction between the two. Regression coefficients were tested using a one-sample t-test against 0 (TS: dopamine response size p=0.03, perceptual uncertainty p=0.01, interaction p=0.04. VS: dopamine response size p=0.86, perceptual uncertainty p=0.01, interaction p=0.94). For plots G-I n=6 mice for the VS and n=4 mice for the TS.

Other theories on the role of movement-related dopamine suggest these signals facilitate movement initiation and/or modulate ongoing action (da Silva et al., 2018; Howard et al., 2017; Howe and Dombeck, 2016). To test if TS dopamine release can also support this function, we optogenetically stimulated TS dopamine release, in a close loop manner, when mice were immobile in an open field arena (Figure S6C). Optogenetic stimulation did not influence whether the animal was more likely to move, nor modulate the movement parameters if mice did move (Figure S6D-I). Taken together, these results suggest that movement-related dopamine activity in the TS can act to reinforce state-action associations but not influence ongoing action.

As stimulation of dopamine in the TS biased choice on the subsequent trial, we sought to determine whether endogenous dopamine release in the TS acted in the same way to bias behavior during the frequency-discrimination task. To do this we classified trials based on whether the recorded TS dopamine signal was large or small (Lak et al., 2020) (see methods) and examined how this affected the choice bias on the subsequent trial. We found that there was a strong relationship between the magnitude of the TS dopamine response and the choice bias on the next trial, specifically that mice were more likely to repeat the action they had taken on the previous trial if there had been a large dopamine response in the TS on the previous trial (Figure 6G, 6H and 6J). This relationship between the dopamine response on the previous trial and the subsequent choice was related to the degree of uncertainty on the current trial, with mice exhibiting a stronger bias when there was a high degree of sensory uncertainty for the current trial (Figure 6H). In contrast, we found no relationship between the size of the dopamine reward response in the VS and the choice bias on the subsequent trial (Figure 6I and 6J). Taken together, these results show that movement-related dopamine release at the time of choice acts as a value-free teaching signal to reinforce stimulus-action associations in the TS, such that mice learn to repeat the action they have taken in the past when they hear the auditory stimulus.

### A dual-controller model recapitulates the effects of experimental manipulations

To understand the interactions between the striatal regions that receive reward or action prediction errors, we built a basal ganglia model where we could simulate learning in regions that were updated by APE or RPE (Figure 7A). Our model consisted of three sets of striatal neurons that received auditory state information and could learn to control actions and generate predictions for the calculation of the APE and RPE. The value-based control system where learning was driven by RPE, consisted of an actor and a critic striatal region that learned to control actions and generated reward prediction, respectively. The value-free control system that mimicked the TS, was updated by APE and learned to control action and generate action predictions. We first aimed to understand how each system learns independently and in conjunction. We found that a model with only a value-free controller, which relied exclusively on updates by APEs, was unable to learn the task. This was expected as the model continues to repeat what it has done in the past, which is to choose left or right at random in equal proportion, so no particular stimulus-action association is preferentially strengthened in response to each sound (Figure 7A). In contrast, a model with only the value-based controller, which is updated by RPEs, was able to independently learn the task as these prediction errors update the value of actions in a particular state (action values). Due to this the network learns over time to select the actions that maximize the cumulative reward (Figure 7B). Interestingly, a model with dual-control systems, updated by reward and action prediction errors, learns quicker than the single RPE learning system (Figure 7B). Initially, there is no difference between the learning rate of the two models, because early learning is exclusively driven by RPE updates, for the same reasons explained above. However, as the dual model begins to consistently choose a particular action in response to a particular sound the APE updates start to preferentially reinforce specific stimulus-action associations. These associations then contribute to driving choices and in turn provide a boost to the performance and the learning rate. This pattern of divergence between the value-based and dual-controller model recapitulates the learning deficits we observed when we either lesioned the TS (Figure 1I) or removed the dopamine innervation to this region (Figure 2B). In both cases there was no difference between the groups initially and then the control groups (i.e., intact dual control system) increase their learning rate once mice are performing the same action in response to the same sound (Figure S7A). Taken together our model shows that the value-free controller alone cannot support learning but when paired with the value-based controller, it acts to boost learning and store the stable associations.

**Figure 7:**
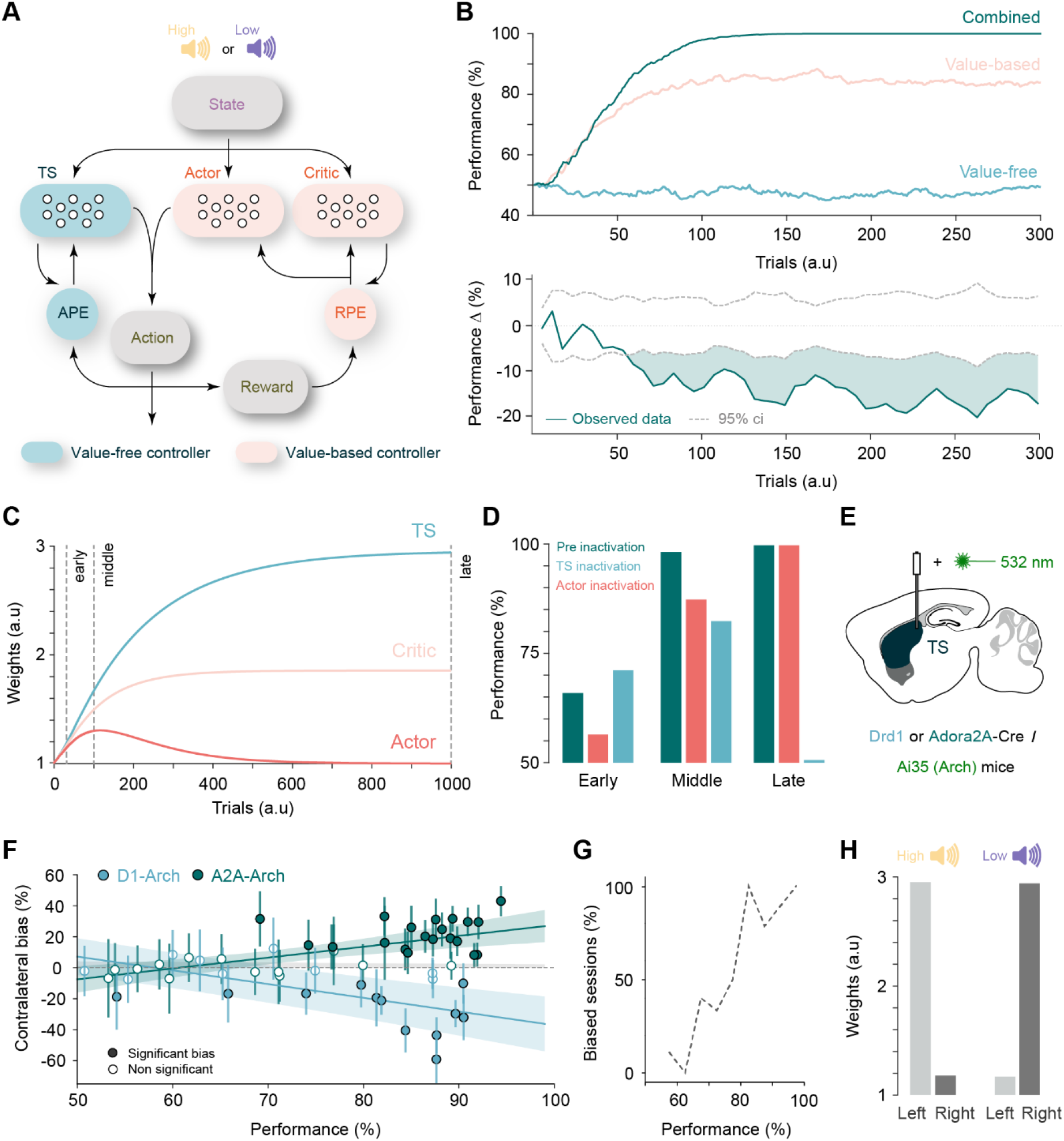
A dual-controller model predicts the effect of experimental manipulations. **A** - Schematic of the network model. **B** - Top graph: Task performance across learning for the full dual-controller model (combined), just the value-based controller, or just the value-free controller. Bottom graph: differences in performance between the combined model and the value-based model (12 random agents selected for each). Dotted lines indicate the 95% confidence interval for the shuffled data (see methods). **C** - Change of the model weights for a reward association (High tone->left action), as means for 100 agents. Vertical lines indicate inactivation time points in D. **D** - Performance levels before and after (as the mean after 10 trials) model inactivations of the TS or the actor networks. **E** - Schematic of the experimental approach for acute inhibition of D1 SPNs or D2 SPNs in the TS. **F** - Quantification of the contralateral bias on opto-stimulated trials (see methods) for each session as a function of session’s performance. Error bars represent standard deviation. Lines show the mean and standard deviation of linear fits for each mouse in each dataset (D1 Arch p=0.004; A2A Arch p=7.6×10^−4^; see methods). **G** - Proportion of significantly biased sessions as a function of performance. **H** - Final weights in the TS for each sound-action association.

Although both the value-based and value-free controllers learn to drive behavior, our muscimol inactivation data as well as previous results (Guo et al., 2018) (Figure 1C), highlight that in well trained mice behavioral performance relies exclusively on the TS. Because the value-free controller cannot learn the task alone, this suggests that behavioral control must be transferred over time from the value-based to value-free controller. In the previous models that proposed a value-free learning system the interaction between this, and the value-based system was decided by an additional arbitration process (Bogacz, 2020; Miller et al., 2019). In our model we sought to explain the transfer from value-based to value-free control with only the known basal ganglia circuitry, as the biological evidence for an arbiter between the VS and TS is scarce. In our model, the synaptic weights in both the critic and the TS were stably retained but the weights in the actor were allowed to decay over time. The result of this is that as the dual-controller model begins to learn, the synaptic weights in actor, critic and TS regions are all updated (Figure 7C). However, as the model reaches maximal performance on the task the weights in the actor begin to decay because there is no longer any RPE to continue to update and maintain the weights (Figure 7C and S7B). Therefore, what the dual-controller model retains is the predicted value of each state-outcome association in the critic and the state-action association in the value-free controller (TS). The behavioral consequence of this is that early in training the model performance is dependent on what has been learnt in the actor (Figure 7D, left). However, as learning progresses the appropriate state-action associations begin to form in the value-free controller and start to contribute to behavioral performance (Figure 7D, middle). Late in learning, as the synaptic weights in the actor decay, the behavioral performance comes to rely, as with our experimental data, exclusively on the value-free controller (Figure 7D, right).

Our dual-controller model predicts that initial learning is driven in a value-based manner and then rapidly consolidated and stored in the value-free system. If the TS indeed corresponds to the value-free system, it should only come to influence behavior later during training. To explicitly test this, we optogenetically inactivated either types of TS projection neurons (D1 or D2 SPNs) throughout learning (Figure 7E). In line with the prediction, early in training optogenetic inactivation of neither D1 nor D2 SPNs in the TS had a significant effect on behavior (Figure 7F). However, as training progressed the inactivation began to have a significant effect on task performance and these effects grew larger as the mice became experts at the task. These behavioral effects began to consistently increase after the mice were performing above 65% (Figure 7F and 7G). This is a comparable training stage to where the control and lesion groups diverged when the TS was lesioned experimentally or in the model (Figure 1I, 2B and 7B). Taken together, this data shows that the TS only begins to contribute to task performance and learning once mice are already preferentially choosing a specific action in response to the auditory cues.

In the “cloud-of-tones” task, frequency specific corticostriatal plasticity that can support the appropriate sound-action association develops in each hemisphere of the TS throughout learning (Ghosh and Zador, 2020; Xiong et al., 2015). To determine if APE updates are sufficient to establish this frequency specific plasticity we examined the strength of sound-action association weights that form in each “hemisphere” of the value-free controller. In line with experimental data, the appropriate state-action associations form in each hemisphere of the value-free half of the dual controller model, where the updates in synaptic plasticity are exclusively driven by APEs (Figure 7H). Taken together, our behavioral results, chronic lesions, acute inactivation, dopamine recordings, optogenetic perturbations and previous synaptic plasticity results (Xiong et al., 2015) are consistent with the TS forming the value-free half of a dual value-based/value-free learning system.

## Discussion

### Summary

Here we show that movement-related dopaminergic activity in the TS acts as a teaching signal to reinforce state-action associations. The TS dopamine activity is consistent with encoding an action prediction error, the difference between the action that is taken and the extent to which an action was predicted in a given state. This value-free signal serves to teach mice to repeat the actions that have been taken in the past. Alone the value-free system (APE->TS) is not able to support reward-guided learning but in conjunction with the canonical reward prediction error system it learns to mimic and store the value-guided state-action associations. We show that when combined with the value-based, RPE-updated basal ganglia circuitry, the value-free control system acts to boost and consolidate learning about consistent state-action associations. Together we show that there are two types of dopaminergic prediction errors that work in tandem to support learning, each reinforcing different types of association in different areas of the striatum.

### All movement-related dopamine signals may encode action prediction errors

The identification of dopamine transients prior to movement initiation led to the idea that movement-related dopamine release may trigger the initiation of action (da Silva et al., 2018; Howard et al., 2017; Howe and Dombeck, 2016; Klaus et al., 2019). Initial experiments supported this idea as optogenetic stimulation of the SNc dopamine release at supraphysiological levels triggered the initiation of movement and increased movement vigor (da Silva et al., 2018; Howard et al., 2017; Howe and Dombeck, 2016). However, in line with our results, more recent experiments that used optogenetic stimulation parameters that were calibrated to mimic physiological levels of SNc dopamine release did not trigger body movement or lead to an invigoration of ongoing action (Coddington and Dudman, 2018). The calibrated stimulation parameters were however, as we observe, able to act as a teaching signal to drive conditioned place preference learning. In line with this, optogenetically stimulating the SNc, where movement-related dopamine activity is prominent, reinforces stimulus-response associations (Saunders et al., 2018). In contrast, optogenetic stimulation of the VTA, where reward prediction error signals (RPE) dominate, supports stimulus-outcome associations (Saunders et al., 2018). This suggests that, as we have seen in the TS and consistent with APE, movement-related dopamine activity in other striatal areas may also act as value-free teaching signal rather than as a trigger or modulator of ongoing movement.

In theory for APE to function as a teaching signal it need only reflect scalar information about action and the degree to which actions are predicted (Bogacz, 2020; Lindsey and Litwin-Kumar, 2022; Miller et al., 2019). This is to say the APE term need not be action-specific. In line with this, the activity of SNc movement-related dopaminergic neurons is only broadly tuned to action (da Silva et al., 2018). Our results show that the APE signal is not a pure scalar action signal as the APE response is larger for contralateral rather than ipsilateral actions. This is unsurprising as motor control circuitry is lateralized, with each hemisphere controlling movements on the opposite side of the body. The inputs to dopaminergic neurons in the SNc conform with this pattern as they receive the vast majority of their inputs *from* the ipsilateral hemisphere (Watabe-Uchida et al., 2012). APE may therefore reflect a scalar term with respect to the actions that each hemisphere controls. Finally, there is some evidence that SNc movement-related dopamine activity may also decrease with practice in a lever-pressing task (da Silva et al., 2018), as it should if the activity reflects an APE. Taken together, the current data on movement related dopamine activity is consistent with action prediction error, suggesting that all movement-related dopamine activity throughout the striatum may reflect APEs that serve as value-free teaching signals. Value and movement-related dopaminergic neurons are predominantly arranged along a ventromedial/dorsolateral anatomical gradient (Cox and Witten, 2019). We propose that this reflects a gradient of the two types of dopaminergic teaching signals, RPE coding dominating in the medial ventral tegmental area and APE coding dominating in the lateral portion of the substantia nigra (Figure S7C).

### Relation to other theories

In line with our results, the caudate tail (CDt), primate homologue of the TS, is also innervated by an anatomically distinct population of dopamine neurons (Kim et al., 2014) that are not activated by differences in reward outcome (Kim et al., 2015). Rather, as we observe here, these neurons are preferentially active prior to contralateral orienting eye movement or when stimuli that predict contralateral orienting responses are presented (Kim et al., 2015), as would be expected if they also encode an APE. The contralateral nature of the response in primates was interpreted as a unilaterally processed visual response (Hikosaka et al., 2019; Kim et al., 2015; Kim and Hikosaka, 2015). Our results now show that at least for auditory stimuli, the TS responses in mice are movement and not sensory related. In addition, rabies tracing has revealed that TS-projecting dopaminergic neurons receive input from the deeper motor-related layers of the superior colliculus rather than from the superficial sensory-related layers (Menegas et al., 2015). Taken together we propose that the activity prior to a contralateral orienting eye movement is more likely to reflect an APE rather than as previously proposed a salient stimulus response. Whether APE signals are driven by the preparation or execution of action will need to be elucidated in future work and may help resolve the difference in explanation. While the interpretation of the signal was different, the proposed function of the CDt-projecting dopamine neurons is entirely consistent with what we show here, in that it serves to reinforce stable sensory-motor associations in the CDt in an outcome independent fashion (Kim et al., 2015; Kim and Hikosaka, 2015). This suggests that the TS is unique across species in that it exclusively receives a value-free movement-based teaching signal to update stable sensory-motor associations.

In line with our results, other labs have also shown that dopaminergic activity in the TS is not activated by receipt of reward (Menegas et al., 2018; Menegas et al., 2017). However, the dopaminergic activity is activated by threatening stimuli, such as an air puff or loud sounds. In addition, large TS dopaminergic transients precede avoidance behavior, and these responses decrease as mice stop performing avoidance responses (Menegas et al., 2018). These observations led to the proposal that TS dopaminergic activity encodes a threat prediction error (TPE) (Menegas et al., 2018; Watabe-Uchida and Uchida, 2018). This interpretation is supported by causal manipulations showing that TS dopaminergic activity is needed to reinforce threat avoidance behavior (Menegas et al., 2018). While future work will be needed to determine how TPE can be reconciled with our findings and those from primates that highlight the contribution of this pathway to reinforcing stable sensory-motor associations, we will speculate here on a couple of points. While movement is unlikely to account for all TS dopamine responses, the observations that dopamine activity in the TS is activated prior to avoidance behavior is consistent with our results that TS dopamine activity is activated in response to movement initiation. Indeed, the behavioral reactions to loud sounds or air puff stimuli may also drive part of TS dopamine activity rather than the threat prediction *per se*. While movement may account for some of the observed “threat” responses, it is unlikely to account for them all especially as causal manipulations show that TS dopamine pathway is needed for reinforcing threat avoidance (Menegas et al., 2018). An additional possibility is that TPE and APE are encoded in the activity of genetically different populations of dopamine neurons that project to the TS (La Manno et al., 2016; Poulin et al., 2018; Poulin et al., 2014). Another option is that threat and action are two types of information that excite the same dopamine neurons. In line with this, TS-projecting dopamine neurons receive input from the deep layer superior colliculus (Menegas et al., 2015), which encodes information about both threat and orienting movements (Evans et al., 2018; Hikosaka and Wurtz, 1983; Lee et al., 2020). Taken together it’s clear that dopamine activity in the TS has multiple roles in both learning stable sensory-motor associations as well as in reinforcing behavioral avoidance, which should be noted is another form of sensory-motor association. Future work will be needed to determine if these roles are supported by the same or distinct dopaminergic populations.

### Connection between action prediction errors and habit formation

APE was first conceived as a teaching signal that could update habitual value-free state-actions associations (Bogacz, 2020; Miller et al., 2019). Our results now show that dopamine neurons that project to the TS encode this type of movement-based prediction error. Interestingly the CDt is where habitual stimulus-action associations are stored (Kim et al., 2015; Kim and Hikosaka, 2013; Yamamoto et al., 2013). While further work is needed to test if CDt projecting dopamine neurons also encode APE, it is notable that the dopamine->CDt system was not thought to be able to learn by itself because it is also updated by an outcome-independent teaching signal (Kim et al., 2015; Kim and Hikosaka, 2015). Rather, as we show works in our model, it was proposed that the dopamine->CDt could learn to mimic the actions that have been driven in the past by a “goal-directed” region of the striatum, which in turn learns the value of actions directly (Kim and Hikosaka, 2015). The primate goal-directed region is the caudate head, where neurons, updated by RPEs, rapidly learn the short-term value of actions (Kim and Hikosaka, 2015). As with the *Actor* in our dual-controller model, these short-term associations then decay quickly (Kim and Hikosaka, 2013). Here, we show how such a dual value-based/value-free learning system can be implemented and work to accelerate and consolidate learning. Taken together, we propose that habitual (value-free) and goal-directed (value-based) associations are learnt in different regions of the striatum based on different prediction errors signaled by different dopaminergic populations. RPEs are used to update the short-term value-based associations in the anterior striatum/caudate that guide goal-directed behavior. In contrast, APEs from the SNl update the long-term value-free stable associations in the CDt/TS to guide habitual behavior.

### Advantages of a value-free system

The value-free system updated by APEs learns to repeat the actions that have been taken in the past. This does not seem like an advantageous strategy compared to relying on the value-based system that encodes the relationship between rewards and actions. So why have this system? The first thing to note is that while APE is a value-free teaching signal it does not mean that the associations that are formed in the TS will be insensitive to value. Rather the “value-free” association forms through repetition of actions that were initially taken in pursuit of value. In this way the associations that form in the TS can be thought of as storing the long-term “value” of an association even though they were never reinforced in value-based manner *per se*. The resulting association in the TS will then be sustained in an outcome independent manner so they are stably retained. Indeed, stimulus-action associations in the CDt system are incredibly stable, once formed they are retained for >100 days in the absence of reinforcement (Yasuda et al., 2012). Therefore, the first advantage of a value-free teaching signal is that associations are stored in an outcome-independent fashion such that they are stably retained. Another advantage of the value-free system is that from a normative perspective it reduces the complexity of the policy, i.e., the mapping from states to actions (Gershman, 2020). This reduction in policy complexity theoretically increases the efficiency for storing state-action associations. In line with this, habitual responses are often faster and more efficient (Wood and Runger, 2016). Overall, the value-free system could ensure that consistent associations are stored in a more stable and efficient manner that allows them to be acted upon quickly and reliably.

Another idea from the Bogacz lab is that stimulus-action associations in the value-free system form a prior for Bayesian inference of action that encodes the action policy that has worked in the past (Bogacz, 2020). During the inference this prior would be useful in biasing decisions when there is a high degree of uncertainty regarding the value of the action computed in the value-based controller. Indeed, under time pressure people default to repeating the choices they have previously made (Betsch, 2004) and when cognitive resources are taxed subjects increase their reliance on habits (Wood and Runger, 2016; Wood, 2014).

Recent theoretical work has also predicted that if dopamine neurons encoded an APE, then it would allow the basal ganglia to learn from any area of the brain that controls action (Lindsey and Litwin-Kumar, 2022). In our model the value-free system learns to mimic the actions that were initially driven by the value-based basal ganglia system. This initial control however need not come from the basal ganglia. If another system, such as the motor cortex or cerebellum, which can control action independently from the basal ganglia, is driving actions then the value-free system can equally learn to mimic and repeat the state-action associations that are initially driven by these systems. Once a state-action association is being performed consistently the APE system will store these associations stably no matter why the actions were initially performed. In this way dopamine encoding APE can support learning from experiences that were not driven by the basal ganglia’s policy, in other words APE coding allows “off-policy” learning.

## Conclusion

In conclusion, we have identified that dopamine neurons projecting to the TS encode an action prediction error. This value-free movement-based teaching signal acts to reinforce specific stimulus-action associations in the TS by learning to repeat the actions that have been taken in the past. This movement-based teaching signal ensures that associations are retained stably in an outcome independent manner. Future work will be needed to determine whether all movement related dopamine signals are indeed APEs. In addition, it will be important to establish the exact nature of the interaction between the value-based and value-free basal ganglia systems and to elucidate the contribution of this teaching signal to habits, priors, and off-policy learning.

## Materials and Methods

### Animals

Male and female adult mice from the following mouse lines were used: C57BL/6J wild-type Drd1-Cre (Gensat: EY262) and Adora2a-Cre (Gensat: KG139), DAT-Cre (JAX Stock No: 006660), Ai14 (tdTomato, JAX Stock No: 007914), Ai35 (Arch-GFP, JAX Stock No: 012735) and Ai32 (channelrhodopsin-2/EYFP, JAX Stock No: 024109)

Mice were housed in HVC Cages with free access to chow and water on a 12:12 h inverted light:dark cycle and tested during the dark phase. For behavioral experiments, mice were water deprived. Animals had access to water during each training session, and otherwise 1mL of water per mouse was administered. Water was supplemented as needed if the weight of the mouse was below 85%. All experiments were performed in accordance with the UK Home Office regulations Animal (Scientific Procedures) Act 1986 and the Animal Welfare and Ethical Review Body (AWERB). Animals in test and control groups were littermates and randomly selected.

### Behavioral procedures

#### -Training box

Mouse training was carried out in a sound-isolated box containing the behavioral chamber, which consists of a closed arena of 19 by 15 cm with three ports located in one wall (1.95cm from the floor, each port was located 3cm from each other). To better separate cues and actions and to increase variability in movement times, the three ports were spaced further apart for photometry experiments. The center port was located in the middle of one of the long sides of the arena and the choice ports were located in the center of the shorter sides (this can be seen in Figure S4 G). The ports are equipped with LEDs so they can be illuminated, and photovoltaic infrared sensors to detect poke events. Water can be delivered through each of the side ports. Ports were purchased from Sanworks or constructed in-house to the same standards. Controlling chips were purchased from Sanworks. The chamber was illuminated exclusively with infrared light. Mice were monitored through standard webcams without the infrared filter. Sounds were delivered through one central speaker (DigiKey part number: HPD-40N16PET00-32-ND) and amplifiers (DigiKey part number: 668-1621-ND). Bpod (Sanworks) was used to control the state machine, and the software was run using Matlab.

#### Cloud of tones task

The cloud of tones (CoT) task is a self-initiated 2-alternative-choice paradigm. The possibility to start a new trial is signaled by turning on the center port LED. Mice start a trial by poking in the center port and holding their position for 100 to 300 milliseconds. The center port LED is turned off after that required time has elapsed. In between 0 and 50 milliseconds after poking in the center port, sound is triggered, lasting for 500ms. Following poking out from the center port, mice were trained to poke in either of the two side ports. 2uL of water is delivered in only one of the two ports, contingent on the stimulus.

#### Training procedure

Depending on the training stage, pokes in the wrong port abort the trial. During the first day of training, mice were habituated to the box and the poking sequence by doing a visual version of the task in which both side ports delivered water in every trial. Water amounts were decreased from 5uL to 2uL during the first 3 days of training. All photometry recordings started after the initial habituation day and after the reward amount was decreased, but before the animals’ performance improved beyond chance.

For the simple version of the task (not the psychometric), we included an anti-bias protocol to force the animal to sample from the two ports. The protocol samples every ten trials for the proportion of error trials and calculates the percentages of port choices on those trials, adjusting the target port on the next ten trials to overcome the bias, proportionally to the errors and the bias. Therefore, this protocol engages progressively more as the animal becomes more biased and disengages if the animal becomes unbiased. This protocol was only active during the initial phases of learning as it uses the proportion of error trials for engagement and calibration. All the experiments using the psychometric version of the task were performed once mice were experts (>85% performance on the simple task). The simple version of the task consisted of delivering only the easiest possible trial types, with 98% of tones being from one of the two octaves (see below). For the psychometric experiments, 7 equally spaced ratios of high or low tones were used. From the beginning of training, stimulus type was randomly interleaved, affected by the anti-bias protocol described above.

#### Variations of cloud of tones task

Several variations of the cloud of tones task were used during photometry recordings. In the outcome value change experiment, unexpected large rewards (6 ml) and omissions were introduced randomly on both sides with a probability of 0.1 each. Mice experience multiple sessions of this protocol to get enough trials of each type.

In the predicted value change experiment, the value of one port was changed 100 trials into a session. This port delivered 6 ml rather than 2 ml, whilst the other port still delivered 2 ml. Mice did multiple sessions of this protocol and experienced both sides becoming the large reward size.

The state change experiment was performed only once per mouse. 150 trials into a training session, the sound that had previously corresponded to the contralateral choice to the recording fiber was changed for a white noise stimulus. This new white noise stimulus overlapped with both original cloud of tone frequencies. This white noise was louder than the background white noise and was clearly detectable. The mice were required to make a contralateral response to the white noise to receive a reward.

To test whether the dopamine signals were dependent on an auditory stimulus being present, trained animals were tested by the sound for which the correct choice would be contralateral to the recording site being omitted.

#### Sound stimuli

Sounds consist of a stream of 30ms pure tones presented at 100Hz (each tone was introduced 10ms after the previous one). One of two octaves (5 to 10KHz, or 20 to 40KHz) was selected as the target octave to indicate the side port where water was available on that trial. Each 30ms tone was randomly drawn from 16 logarithmically spaced frequencies on each octave. The difficulty of the trial was controlled by varying the proportion of 30ms tones from each octave. For example, in a trial cataloged as 82% of high tones, for each 30ms tone there are 82% chances of playing a tone from the 20-40KHz octave, and 18% (100 - 82) chances of playing a tone from the 5-10KHz octave. These probabilities are independently computed, so two tones from different octaves can sound simultaneously. The overall amplitude of the sound is randomly selected between 60 and 80 dB.

#### Water delivery

Water was delivered into the two side ports for correct choices using a solenoid valve that was carefully calibrated. As previous studies have suggested that the neurons in the TS are responsive to valve click noises (Menegas et al., 2017), for the photometry experiments we placed valves outside the training boxes and muffled them using sound insulation foam. In addition, we played quiet white noise constantly inside the training box. We confirmed with a microphone that the valve clicks could not be detected with these precautions.

#### Open-field stimulations

For this experiment we used a 40cm by 40cm square arena, and experiments were conducted with ambient light. Sessions lasted for 30 minutes. See *optogenetic manipulations* for specifics about the triggered stimulations.

#### Open-field photometry recordings

For this experiment we used a 50 cm (l) by 20cm(w) by 28 cm (h) arena. Animals were allowed to explore the setup for 20 minutes, during which video footage and photometry recordings were acquired.

### Surgical procedures

Mice were anesthetized with Isoflurane (0.5–2.5% in oxygen, 1 l/min), also used to maintain anesthesia. Carpofen (5 mg/kg) was administered subcutaneously before the procedure. Craniotomies were made using a 1-mm dental drill (Meisinger, HP 310 104 001 001 004). Injections were delivered using pulled glass pipettes (Drummond, 3.5”) in an injection system coupled to a hydraulic micromanipulator (Drummond, Nanoject III) on a stereotaxic frame (Leica, Angle Two™), at approximately 10 nl min^−1^. For optogenetic experiments, we used flat optical fibers of 200μm diameter (Newdoon: FOC-C-200-1.25-0.37-7) and tapered optical fibers (Optogenix, Lambda Fiber Stubs) of 200μm diameter with 1.5mm active length, 4mm implant length. Implants were affixed using light-cured dental cement (3m Espe Relyx U200) and the wound sutured (6-0, Vicryl Rapide) or glued (Vetbond). Coordinates are measured from the extrapolated intersection of the straight segments of the coronal sutures between the parietal and the frontal bones. This point usually lies slightly frontal to bregma and is more stereotypic than bregma itself with respect to the brain. Coordinates used for targeting were, TS: 1.8 posterior; 3.45-3.55 lateral; 3.4-3.5 depth, VS: 1.0-1.1 anterior; 1.65 lateral; 4.4-4.5 depth.

### Surgeries for fiber photometry recordings

Photometry fibers were targeted 0.1-0.15 mm above the injection site for both TS and VS. For the TS injection, volumes were between 30 and 100 nl were used. For the VS 90 nl injection volumes were used. In the same surgery, fiber cannulae (Doric Lenses 0.57NA, 400 mm diameter, 4 or 7mm long) were implanted between 0.1 and 0.15mm above the injection site(s). For the cohort of mice with fibers in the TS, 6 had fibers on the left and 7 had fibers on the right. 4 of the VS implants were on the left and 3 were on the right. 8 mice that were targeted for TS recordings were not used in any experiments as initial investigation revealed they had reward responses which have not been observed in the TS (Menegas et al., 2018; Menegas et al., 2017). To confirm these fibers were outside the TS in 6/8 of these mice we performed serial two-photon microscopy and confirmed that in all cases the fibers were located outside of the TS.

### Viral injections

All viruses are made in-house except for pAAV5-CAG-dLight1.1 (Addgene, 111067) which was used for the **photometry recordings**.

**Chronic lesions** of the TS were achieved by injecting a 1:1 mix of AAV2/1-hSyn-Cre 10^14^ vg/ml and AAV2/5-EF1a-DIO-taCasp3-T2A-TEVp 10^14^ vg/ml. The mix was diluted 5 times in saline buffer prior to injection. The same surgical procedure, using the virus AAV2/5-CAG-EGFP (3×10^12^ vg/ml), was used for the control group. In each hemisphere 4 injections (30nl each) at 4-5 different depths were made to distribute the viruses as evenly as possible and to provide enough coverage.

### Pharmacological manipulations

For **muscimol** injections, bilateral cranial openings were performed over the DMS and the TS, and a headbar was positioned stereotactically. A landmark that aligned to bregma allowed for future injections in stereotactic positions. Closures of cranial openings were prevented by covering the exposed brain with Duragel (Cambridge NeurotechⓇ).Skullw ascovered w ith kw iksil(W orld Precision Instruments), which was removed before every injection. Before each training session, animals were headfixed while awake and 30μL of either muscimol (Sigma-Aldrich) at 0.2mg/ml or saline buffer were injected for experimental and control sessions, respectively, co-injected with CTB (Cambridge bioscience) of different colors to trace injection sites during histology. After a 15 minute period, animals started the training session.

For **dopamine neuron ablations** we followed the protocol in (Menegas et al., 2018). Briefly, we first injected intraperitoneally (10mg/kg), a solution made of 28.5 mg desipramine (Sigma-Aldrich, D3900-1G), 6.2 mg pargyline (Sigma-Aldrich, P8013-500MG), 10 mL water, and NaOH to pH 7.4. Subsequently animals underwent surgery (see above), and a solution of 10 mg/mL 6-hydroxydopamine (6-OHDA; Sigma-Aldrich, H116-5MG), dissolved in saline buffer was injected. Once prepared, we kept the solution on ice covered from light, and injected it within 3 hours. If the solution turned brown it was discarded. For controls we only injected saline buffer.

### Optogenetic manipulations

For **opto-inhibition** of either the D1 and the D2 pathways, in 15-25% of trials, randomly selected, a sustained pulse of green (532nm) light was delivered after mice initiated the trial, always preceding the onset of sound delivery by at least 50ms and lasting more than the sound duration. Light intensity was calibrated to 12mW at the fiber tip.

For **dopamine opto-excitation**, during the CoT task, the first 150 trials of each session were done without stimulation to get a baseline behavior, and subsequently, stimulated trials were introduced with these parameters: blue (473nm) light delivered starting at the time of poking and lasting 150ms in 5ms pulses at 33Hz of 4mW intensity measured at the fiber tip. For the state-action experiment, stimulation was delivered unilaterally in the center port for trials in which the state predicted a movement contralateral to the stimulated hemisphere. For the state-outcome experiment, stimulation was delivered bilaterally on one of the two side ports (for the whole duration of the session) every time the mice chose that port (correct and incorrect trials), coupled with water during correct trials. No anti-bias protocol was employed during these experiments. To test for effects on movement initiation, we performed an experiment similar to (da Silva et al., 2018). We placed mice in an open field where mice received a blue (473nm) light stimulation (5ms pulses at 33Hz during 500ms of 4mW intensity measured at the fiber tip) if they were immobile for at least 1 second. For each of these events, there was a 50% chance of triggering the laser. Trials in which light was not delivered were used as within-animal control.

### Immunohistochemistry

Brain slices were all stained following the same procedure: Blocking in staining solution (PBS + 1%BSA + 0.5%Triton-X) for 15 minutes. Primary antibody(s) (1:500 in staining solution) for 2-4 hours at room temperature or overnight at 4 degrees celsius while rocking. 15 minutes wash with staining solution. Second antibody(s) (1:500 in staining solution) and DAPI for 2 hours at room temperature while rocking. Slices were then washed in PBS and mounted using Mount Medium. Primary antibodies used were NeuN, TH and GFP (to reveal dLight-expressing cells). Secondary antibodies used were Alexa-488 anti-mouse, Alexa-567 anti-chicken, and Alexa-647 anti-rabbit.

### Tissue processing and image analysis

Animals were perfused with PBS followed by 4% PFA, and brains were extracted and fixed in 4% PFA for 24 hours and incubated in PBS.

#### Muscimol injection and fiber placement locations

We imaged the brains using serial section (Mayerich et al., 2008) two-photon (Ragan et al., 2012). Our microscope was controlled by ScanImage Basic (Vidrio Technologies, USA) using BakingTray, a custom software wrapper for setting up the imaging parameters (https://github.com/SainsburyWellcomeCentre/BakingTray,https://doi.org/10.5281/zenodo.3631609). Images were assembled using StitchIt (https://github.com/SainsburyWellcomeCentre/StitchIt, https://zenodo.org/badge/latestdoi/57851444)). The 3D coordinates of the injections and fiber placements were determined by aligning the brains to the Allen Reference Atlas (Allen Reference Atlas – Mouse Brain. Available from atlas.brain-map.org.) using brainreg (Tyson et al., 2021), and visualized using custom functions and brainrender (Claudi et al., 2021).

#### Quantification of chronic lesions

Brains were sliced using a cryotome at a thickness of 30um. 15 to 20 slices covering the entire striatum at regular intervals were selected for NeuN staining. Slices were mounted in standard glass slides using standard mounting medium, and subsequently imaged in the Slide Scanner (Zeiss) using a 20x objective. Individual slices were registered to the Allen Reference Atlas (Allen Reference Atlas – Mouse Brain. Available from atlas.brain-map.org.) using ABBA (https://github.com/BIOP/ijp-imagetoatlas), and the NeuN channel was thresholded automatically, per slice, based on the intensity levels in the cortex. The coverage of NeuN staining in the striatum was determined for each slice, and the inverse was determined as lesioned area.

#### Quantification of dopamine neuron ablation

Brains were sectioned and mounted as described for the chronic lesions but stained for Tyrosine Hydroxylase to specifically label dopamine cell processes. Slices were imaged using the Axio Scan (Zeiss). Manual regions of interest (ROIs) were drawn for the striatum, the cortex, and the background in each slice. For each slice, the mean intensity in the striatum was normalized to the cortex following background subtraction, and the relative intensity between the striatum and cortex was calculated. For the analysis of the correlation with the performance, data was normalized within mouse (posterior striatum ratio / anterior striatum ratio).

### Photometry and video acquisition

Fluorophores were stimulated using 465 nm and 405 nm LEDs (Thorlabs) of max power 0.2 mW. The 465nm and 405 nm LED amplitudes were modulated using a sinusoid of 211 and 531 Hz respectively. The 405 nm light falls at the isosbestic point for fluorophores of the type used in this study (Robinson et al., 2019). This enabled separation of signal from movement artifacts and bleaching as done by (Lerner et al., 2015). Light was passed from the LED source through optical fibers with NA 0.57, through a commutator (Doric Lenses, FRJ 1×1 PT 0.15) to a patch cord (Doric Lenses, FC-ZF1.25 LAF). A mini cube and photodetector (Doric Lenses) were used to collect the signal. The signal was then passed to a NIDAQ (National Instruments) and recorded and analyzed using custom Python scripts as described in the *statistical analysis* section. Behavior was controlled with a Bpod which sent TTL pulses to the NIDAQ at the start of each trial.

Animals were filmed from above during training and recording sessions using a Basler acA640-750um USB 3.0 camera. Videos were acquired at 30Hz and synchronized with the photometry acquisition using the NIDAQ. For ease of analysis animals were always trained in the same box and cameras were never moved.

### Statistical analysis

#### Behavioral data preprocessing

Sessions with less than 60 trials were omitted, the first 5 trials of each session were discarded, and trials in which the animal was not engaged (defined as having an inter-trial interval longer than three times the median value of that session) were not considered for analysis. Together, this amounts to less than 2% of data discarded. The remainder of the trials were ordered chronologically. Animals which did not learn the task (end performance less than 55%) were discarded from the analysis. This amounts to a total of 3 animals in the whole study.

#### Learning rate experiments

The first 5000 trials for each mouse were used for analysis. To calculate individual learning parameters, per mouse, we modeled the performance of every mouse using the sigmoid function: (max_performance - 0.5) / (1 + np.exp(-slope * (x - bias))) + 0.5. Data was scaled before the fitting and parameters were rescaled afterwards. The maximum performance was defined as the maximum of the median of the trials, binned using a window of size 200. Parameters *slope* and *bias* were calculated using scipy in python (optimize.minimize function). The maximum learning rate is calculated from the model. Statistical differences between the groups for these parameters were calculated using the non-parametric test Kruskal-Wallis from scipy.stats.kruskal. Significance of the behavioral correlation with the lesion size was performed using the scipy.stats.linregress function. To calculate the differences in performance at different times in learning, we first removed those trials in which mice were extremely biased towards one of the two ports. Bias was determined as described in the *behavioral procedures* section, and extremely biased trials were defined as those having a value larger than twice the standard deviation for the whole dataset. Two mice (one experimental and one control) were removed for the chronic lesions experiment as they did not learn the task at all, and we suspected they were deaf. Additionally, the two last sessions of one control and one experimental mouse were removed as the performance dropped to chance. One experimental animal was excluded from the dopamine cell ablation experiment as the lesion quantification showed no ablation (this mouse performance was comparable to controls). At each point in training, performance was defined as the performance of the past 100 trials. To assess the significance of the differences between the two groups, the data was binned using a window of 100 trials, and the differences between the means of each group were calculated. To generate a shuffled dataset, experimental labels (e.g., lesion or control) were randomly assigned to each mouse, always maintaining the proportion of labels on the original dataset, and differences between groups were calculated the same way. We did this 10,000 times. The same procedure was used to analyze the data from the dual-model controller comparisons. The global significance of the dataset was assessed as the likelihood of the cohorts being different at any time, in comparison to the shuffled groups. The mixed Anova was used from the pingouin package.

#### Opto-inhibition experiments

To assess the significance of the biases caused by the optogenetic manipulations, we generated, for every session, a baseline distribution of port choice proportions for every unstimulated trial type (proportion of high vs low tones), using the same number of trials that were stimulated for that trial type. This generated the natural variability in the potential choices for each trial type and was used to assess the significance of the biases for individual sessions. The total bias for each session was defined as the average difference between opto-stimulated and unstimulated trials for all trial types. Only sessions with more than 150 trials in total were selected for analysis. For animals in which more than data for more than one session and stimulation type was available, we selected the one with the best performance. The statistical significance for each group was calculated using the non-parametric test Kruskal-Wallis from scipy.stats.kruskal, comparing the observed biased values of every session against a randomly-sampled counterpart, per session, using the variance described above. Only sessions in which the animals did the psychometric version of the task were included in this analysis.

#### Dopamine photometry pre-processing

The raw data were demodulated offline using custom Python scripts to produce traces that corresponded to the signal (465nm) and background (405nm) channels. Then the data were processed according to the methods described in (Akam and Walton, 2019). Briefly, the demodulated traces were denoised using a median filter and a low pass Butterworth filter (10Hz cut off). The resulting signal and background channel traces were high-pass filtered at 0.001Hz to correct for photo-bleaching. To correct for motion artifacts, the background channel data was fitted to the signal channel data using a linear regression. The proportion of signal that was explained by the background channel was then subtracted from the signal channel component, such that only the signal specific to the 465 nm excitation frequency remained. Finally, dF/F was calculated by dividing this signal by the baseline fluorescence (the signal channel trace filtered using a low pass filter with a cut off of at 0.001Hz). All traces were z-scored to allow better comparison across animals and sessions.

Peak dopamine responses were calculated using the Python package PeakUtils or by taking the maximum value of the trace if no peak was found in a given window. For cue responses, the window was between the cue onset and entry into the choice port. For action (APE) responses, the window was between the time of exiting the center port and entry into the choice port. For outcome responses (used in the outcome value manipulation experiment) the window was between the time entry into the choice port and 200 ms later. As VS dopamine clearly dips to omitted rewards, instead of calculating peaks, the mean of the dopamine response within this window was used as an estimate of the response.

#### Kernel regression model of photometry signal

Following (Parker et al., 2016), we built a linear regression model to predict the photometry signal at each time point from the behavioral events around this time. In this model, the predicted dLight response is calculated as the convolution of a time series *a, b*, etc. representing different behavioral events as series of 0s and 1s, where 1s represent the occurrence of the behavioral event. This means that at time *t*, the predicted dLight response *g(t)* is given by the weighted sum of the different behavioral events shifted in time within a set window. The model can be expressed as follows:

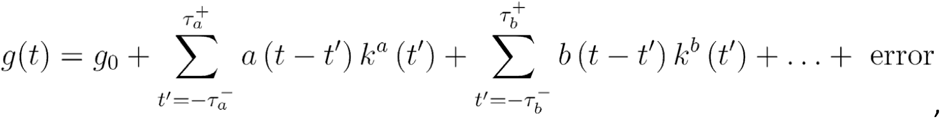

where 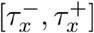 gives the shifted time window in which behavioral event x is allowed to influence the predicted photometry signal. *k*^*a*^, *k*^*b*^ etc. are the kernels for the behavioral events, or equivalently, the linear regression weights for the events at each different time shift. Therefore, they have to be estimated and when plotted as a time series from the estimated response profile for the dLight signal due to a given behavioral event. These regression coefficients were estimated using the linear regression function of the Python package scikit-learn.

Behavioral events were selected from trials in which the animal did not repeat events (for example, did not repeatedly poke their head in the center port). The model was fitted for each mouse for each session as the signal in the TS and VS evolved over learning. As the choice movement initiation time is unclear and may start prior to the withdrawal of the head from the center port is detected, the movement kernels were allowed to extend 0.5 seconds prior to the event. As the movement reaction times were longer than the cue and outcome responses, the movement kernel was allowed to extend 1.5 seconds after the event, whereas the cue and outcome kernel windows were limited to 1 second after the event.

#### Calculation of explained variance by behavioral variable in kernel regression model

The calculation of the percentage variance explained by the full model was performed per session per animal and then averaged across sessions. To calculate the percentage variance explained by each regressor, the predicted dLight signal was recalculated without that particular regressor, inspired by (Tsutsui-Kimura et al., 2020). The explained variance of the new prediction compared to the true signal was then calculated. Finally, the percentage variance explained by the removed regressor was calculated by comparing the explained variance of the full model, *v*_*full*_, to the explained variance when that regressor is removed from the prediction calculation *v*_*partial*_ using the following equation:

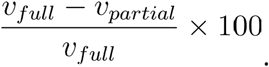

Whilst this method does give a decent comparative measure of the contribution of each regressor to the explained variance of the data by the prediction, it can result in percentages larger than 100 if the predicted dLight signal without that regressor performs worse than the intercept at explaining the data. This can be seen for some of the VS recordings without the outcome regressor.

#### -Video tracking during photometry recordings and quantification of movement parameters

The position of the animal was tracked using DeepLabCut (Mathis et al., 2018) and variables such as speed, acceleration, angular velocity, angular acceleration were calculated using custom scripts in Python. As the videos for the freely moving experiment were taken at a slight angle, the coordinates were transformed into standard space using custom python scripts. Videos taken during the cloud of tones task were taken from above so there was no need for such a transform. Speed and acceleration were calculated using the nose as a marker. For angular velocity and acceleration, a triangle was formed using the nose and the two ears and a line was drawn from nose to the line between the two ear markers. This line from the nose to the back of the head was taken as the animal’s heading direction. Turn angles were calculated using the cumulative angular velocity. In the task, 0 degrees was defined as the angle when the animal leaves the center port. To calculate the maximum turn angle in the task, a sigmoid curve was fitted to the cumulative angular velocity for each trial between the sound onset and entering the choice port. The upper plateau of the sigmoid (maximum fitted cumulative angular velocity) was then taken to be the turn angle for the trial.

To determine the coarse relationship between the turn angle and the size of the dopamine response, trials were divided into 4 quantiles based on the size of the dopamine response between the animal leaving the center port and entering the choice port for contralateral choices. Turn angles were divided into using the quantiles that were created from the size of the dopamine response. A regression slope was fitted (using the function stats.linregress from the Python package SciPy) between the average quantile turn angle and average quantile dopamine response for each mouse for each session. The fit slope was then compared to the fit slope if the x and y quantile labels were shuffled in. The data was shuffled 100 times and the distribution of the fit slopes from the actual data was compared to the shuffled distribution.

In the freely moving experiment, turns were defined as the moment when the angular velocity crossed a threshold of ± 0.5 standard deviations. Turns were required to last at least 0.06 s and were only included for analysis if no other turns occurred in the preceding or subsequent 0.5 s. The head angle at the times of turn onset were then used to define 0 degrees turn angle and turn angles were determined using the cumulative angular velocity from this time point.

#### Dopamine opto-stimulation experiments

To quantify the bias for each session, we calculated the choice differences between the first 150 trials (pre-stimulation), and the trials beyond 225 (post-stimulation and post-sampling), as long as these were more than 100 trials. Sessions in which the initial bias of the mouse to either of the ports was larger than 2:1 were removed from the analysis. We averaged the biases if several sessions existed for the same mouse on the same stimulation conditions, so that we had only one observation per condition. The random datasets were created by shuffling the trial indexes. We used the non-parametric one-sided Wilcoxon rank-sum statistic (scipy.stats.ranksums) to calculate the significance of the observed biases.

For the open-field stimulation, we calculated the speed of the mouse as the difference between the position of the mouse in each frame, convolved with a kernel of size 5 for smoothing purposes. The overall threshold to consider that the animal was moving was determined empirically, but the results presented were invariant across a large range of values tested. Instant speed was calculated as the average speed from movement initiation to 300ms after. Average movement was calculated from stimulation to 5 seconds later. We used the t-test on two related samples (as above) to assess the significance of every parameter.

#### Analysis of the effect of a large dopamine response on subsequent trial bias

Data analysis was inspired by (Lak et al., 2020). Dopamine responses considered in this analysis were reward responses for the VS and movement-aligned responses for the TS. Dopamine responses were categorized as large or small based on whether they were larger than or smaller than the 65th percentile respectively. The average change in percentage of contralateral choices was calculated for large and small dopamine responses on the preceding trial for each level of sensory uncertainty on the current trial (which correspond to the different percentage mixtures of tone clouds described in *behavioral procedures*). For each mouse a linear regression was performed (using statsmodels.formula.api.ols) to investigate the relationship between dopamine response size on the previous trial, perceptual uncertainty and the response on the current trial. Due to the nonlinear relationship between current perceptual uncertainty and choice bias (Figure 6), a log scale was used for perceptual uncertainty in the regression model. An interaction term between dopamine response size and perceptual uncertainty was also included in the regression model due to the different slopes of the lines in Figure 6H for large and small dopamine. For each term in the regression model, the regression coefficients for each mouse were tested against 0 using a one-sample t-test (scipy.stats.ttest_1samp).

#### Opto-inhibition during learning

For the experiment in which opto-inhibition was done at different points in learning, animals only did the simplest version of the task (stimuli were either 98% of high tones or 98% of low tones). The initial 20 trials of every session were discarded, and the baseline distributions of choice proportions and the session bias were calculated as indicated above. Because mice performed close to perfection in this version of the task, we only included in the analysis the choices to the port where biases were expected to happen, in agreement with our previous results (it is not possible to measure any positive contralateral bias in a trial in which the animal chooses the contralateral port all the time in unstimulated trials). Mice were all implanted bilaterally, and the stimulated implant was alternated every day of training. We only included in the analysis those implant sites which resulted in at least one significantly biased session (p-value < 0.05, calculated as indicated above). This removed 4 implants from the analysis out of 17. The regression models were done for each individual implant, and the average slope for each genotype was calculated. The significance of the global results were assessed against random datasets, generated by shuffles of the performance values associated with every session. P values were calculated as the proportion of ‘random slopes’ being larger (expecting negative and positive contralateral biases for D1-Arch and A2A-Arch, respectively).

## Computational modeling

### Modeling of APE and RPE photometry signal

The RL model used an actor-critic framework with soft-max policy selection. The states and actions available in the task were represented as shown in Figure S5B. The model used a semi-Markov state representation, meaning actor, critic and stimulus-action strengths were updated only if a state transition took place. The reward prediction error was scaled according to dwell time in the state to approximate temporal discounting (Daw et al., 2006). RPE was calculated at each state transition *k* as:

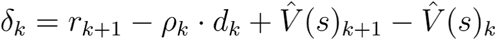

where *r*_*k+1*_ is the reward received in the new state, *ρ*_*k*_ is the average reward per time step and *d*_*k*_ is the time spent in the last state before transitioning. *ρ*_*k*_ was calculated by looking at average reward per time step in the last n (n=500 in our model) state transitions:

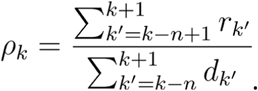

The critic value function was then updated using the RPE ⁳_*k*_ signal:

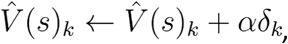

and the actor stimulus-action value function was updated in a similar manner:

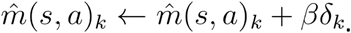

In the above equations, α and β are both learning rates which were both set to 0.005.

As in (Daw et al., 2006), the RPE signal was then taken to be *σ(*⁳_*k*_ *+ ψ)*, where *σ(x) = 0* if *x ≤ 0* and *σ(x) = x* if *x >* 0. *ψ* was set to be 0.1 to represent a baseline level of dopamine.

The non-value dependent model calculated APE, ☐_*ak*_, at state transitions as the difference between the action taken *a*_*k*_ and the stimulus action strengths given the stimulus *A(s)*, according to the equation:

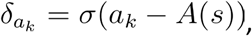

where σ is the rate at which the stimulus-action association was formed, which was set to 0.01 for all simulations apart from the predicted value change experiment simulation, where it was set to 0.001 to mimic how a value-free controller should update slowly. For other simulations this was less relevant as we aimed to show the direction of change of APE and RPE signals rather than their relative rates of updating.

#### Models of novelty, salience, and movement

Every time the animal revisited a state it had visited before, a counter *I(s)* was increased to keep track of how many times that state had been visited:

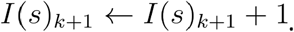

This was used to compute novelty *N*, which decayed exponentially with exposure to a state, according to:

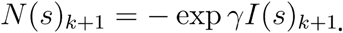

Salience *L* was computed as a weighted combination of value and novelty of the state:

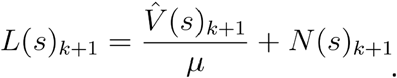

In the above equations, γ and µ were set to 0.01 and 0.5 respectively.

The movement model of dopamine comprised a vector with length equal to the number of actions available. When an action was taken, the corresponding element in the vector was set to 1. All other elements were 0.

In all candidate models of dopamine photometry signals, learning rates and constants were not fitted to match the numbers of trials taken to learn the task seen in the experimental data, so trends should be interpreted as approximations of general patterns rather than models of the exact behavior.

### Dual-controller network model

We simulated a task with two states (s = 1 corresponds to low tone, s=2 corresponds to high tone) and 2 actions (1 corresponding to right and 2 corresponding to left). We modeled two parallel systems, one for the value-based dorsal RPE (separated into an actor and a critic) and one for the value-free tail APE. Each system had a weight matrix that projected from the sensory states to the actions (dimension 2×2),*W*_*actor*_, *W*_*critic*_, *W*_*tail*_. Each system’s output was computed as the weighted sum of the states, i.e.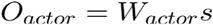 and.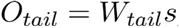

The two outputs were summed up, and an additional noise term was added,*O*_*noise*_ (drawn from a Uniform distribution between 0-1):

. *O*_*total*_ = *O*_*actor*_ + *O*_*tail*_ + *O*_*noise*_ The action associated with the maximal value of *O*_*total*_ was the action that the model as a whole took (corresponding to the animal’s choice). We called it: *a*. The model received a reward, r=1, if the action taken, *a*, corresponded to the action rewarded (here, s =1 to a=1 and s=2 to a=2).

We then computed the RPE and APE signals. For the RPE signal, we first computed the putative action from the critic,

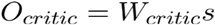. We then computed a putative reward,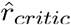, obtained if the maximum of *O*_*critic*_ matched the action rewarded. We computed the predicted reward 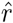 (initialized at 0.5) as a low-pass filter of the reward with a time constant *τ*_*δ*_=100 [a.u.],

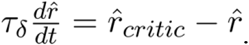

The RPE signal was defined as the reward minus the predicted reward,

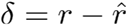

Similarly for the APE, we computed the predicted actions (initialized at 0) as

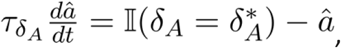

Where 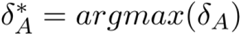 and 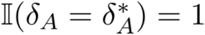 if 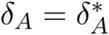 and 0 otherwise.

The APE was calculated as,

*δ*_*A*_=*a−â*, where the time constant 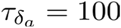 .

Finally, we updated the synaptic weights as a function of the state, the action taken and the APE/RPE signal:

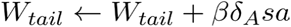, where β = 0.02 is a learning rate,

*W*_*critic*_ *← W*_*critic*_ *+αδsa*, where α = 0.04 is a learning rate,*W*_*critic*_ *← W*_*critic*_ *+αδsa*

The RPE weights of the actor also decayed to its steady-state value of 1 with a time constant of µ=100,

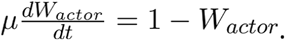

All the weights were bounded to be positive and were initialized to 1.

We simulated 100 trials. A trial lasted 1000 timesteps (2000 for the inactivations). For plotting the performance, we low-pass filtered the reward with a time constant of 10 [a.u.], starting from 0.5. The inactivations were conducted at time 30, 100, 1000 after which either the *O*_*actor*_(for the RPE system) or the *O*_*tail*_ (for the APE system) was set to 0 and weights were not plastic anymore.

## Acknowledgements

We want to thank the Advanced Microscopy Facility in the SWC for assistance with histology processing and image acquisition, especially Jessica Broni-Tabi and Rob Campbell. We are grateful to the Viral Vector Core in the SWC, especially Gemma Estrada Girona and Molly Strom, for the custom production of genetic constructs and adenoviruses. Thanks to Federico Claudi for advice on analysis of the tracking data. We would also like to thank Jesse Geerts for feedback and advice on the APE and RPE dopamine models. We are very grateful to Mitsuko Uchida, Tom Mrsic-Flogel, Sonja Hofer, Rafal Bogacz, Tiago Branco and Andrew McAskill for their comments on the manuscript. Finally, we want to thank all the members of the Stephenson-Jones lab for their valuable contributions during the development of this project.

## Notes

### Competing Interest Statement

The authors have declared no competing interest.

### Summary of Updates

The Author list has been updated

