## Supplemental figures for "Action prediction error: a value-free dopaminergic teaching signal that drives stable learning"

### Supplementary Figures

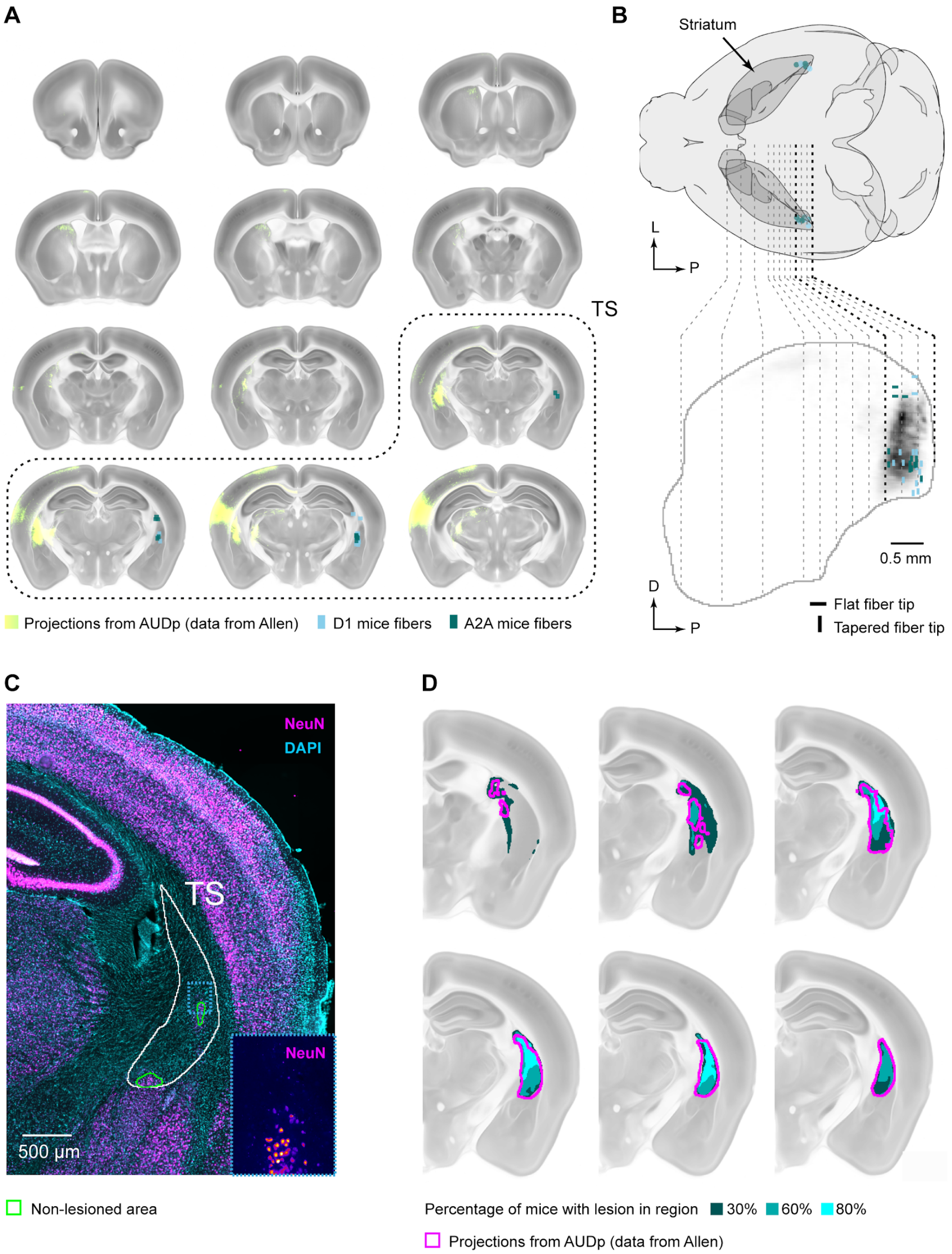


***Figure S1: Histology of optoinhibition and TS lesions. A*** *- Coronal sections along the striatum indicating fiber placement positions (tip of the fiber). Note that fibers were inserted in both hemispheres and are mirrored here for illustration purposes. Primary auditory cortex (AUDp) projections are shown in the other hemisphere.* ***B*** *- 3D rendering (top) and side view (bottom) of the same histological data. On the side view, the striatum is outlined and the AUDp projections are indicated in greyscale.* ***C*** *- Representative image of a lesioned brain illustrating the image analysis used to quantify the proportion of lesioned striatum. Slices were registered to the atlas and the area with remaining neurons (as stained by NeuN) was defined. The rest of the striatum was considered as lesioned.* ***D*** *- Results of the lesion quantification for all the mice in the dataset across several coronal slices of the posterior striatum, that include the entire TS.*

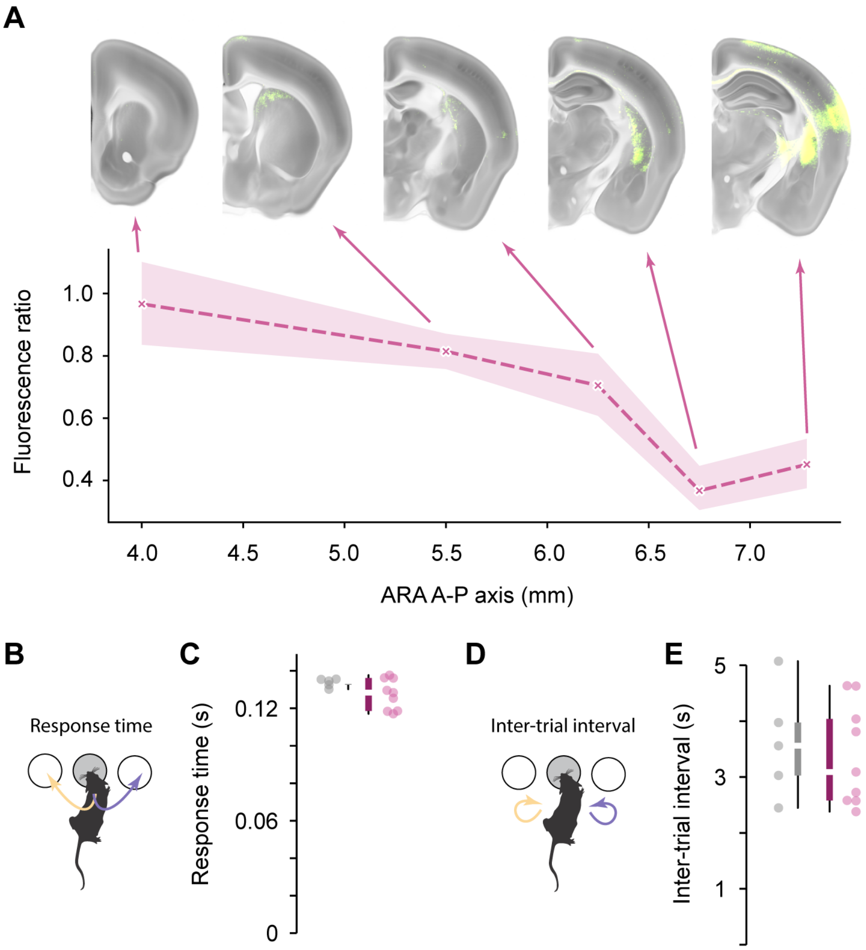


***Figure S2: Histology of TS-dopamine ablated mice and task response parameters. A*** *- Quantification of the TH staining fluorescence ratio between the striatum and the cortex after background subtraction, at different levels in the allen reference (ARA) anterior-posterior axis. The data is shown as the fluorescence relative to controls. Primary auditory cortex projections are shown.* ***B*** *- Time elapsed between center port poke and side port pokes, as medians for each animal, for the 6-OHDA and control groups (p=0.31, Kruskal-Wallis test).* ***C*** *- Time elapsed between trials, as medians for each animal, for the 6-OHDA and control groups (p=0.73, Kruskal-Wallis test).*

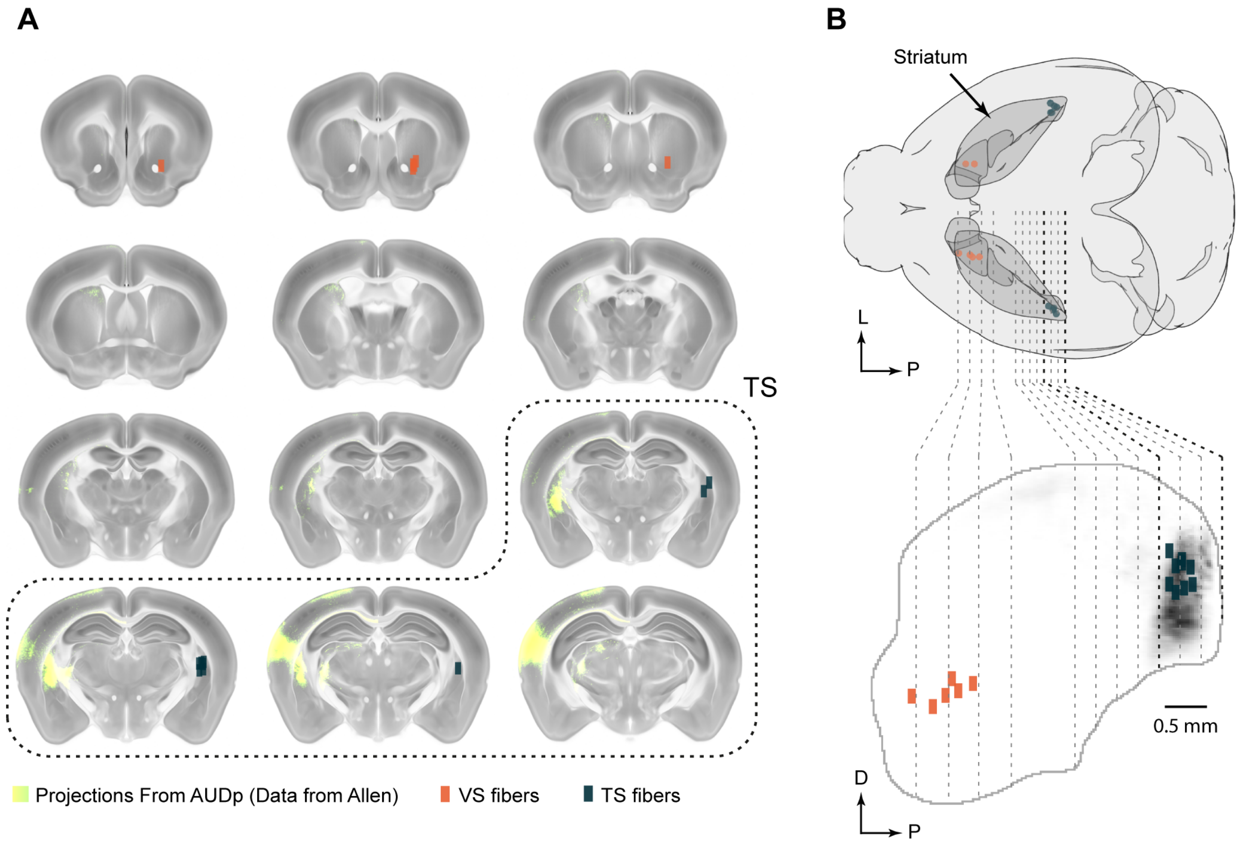


***Figure S3: Histology of fiber photometry placements. A*** *- Coronal sections along the striatum indicating fiber placement positions (center tip of the fiber). Note that fibers were inserted in both hemispheres and are mirrored here for illustration purposes. Primary auditory cortex projections are shown in the other hemisphere.* ***B*** *- 3D rendering (top) and side view (bottom) of the same histological data. On the side view, the striatum is outlined and the AUDp projections are indicated in greyscale.*

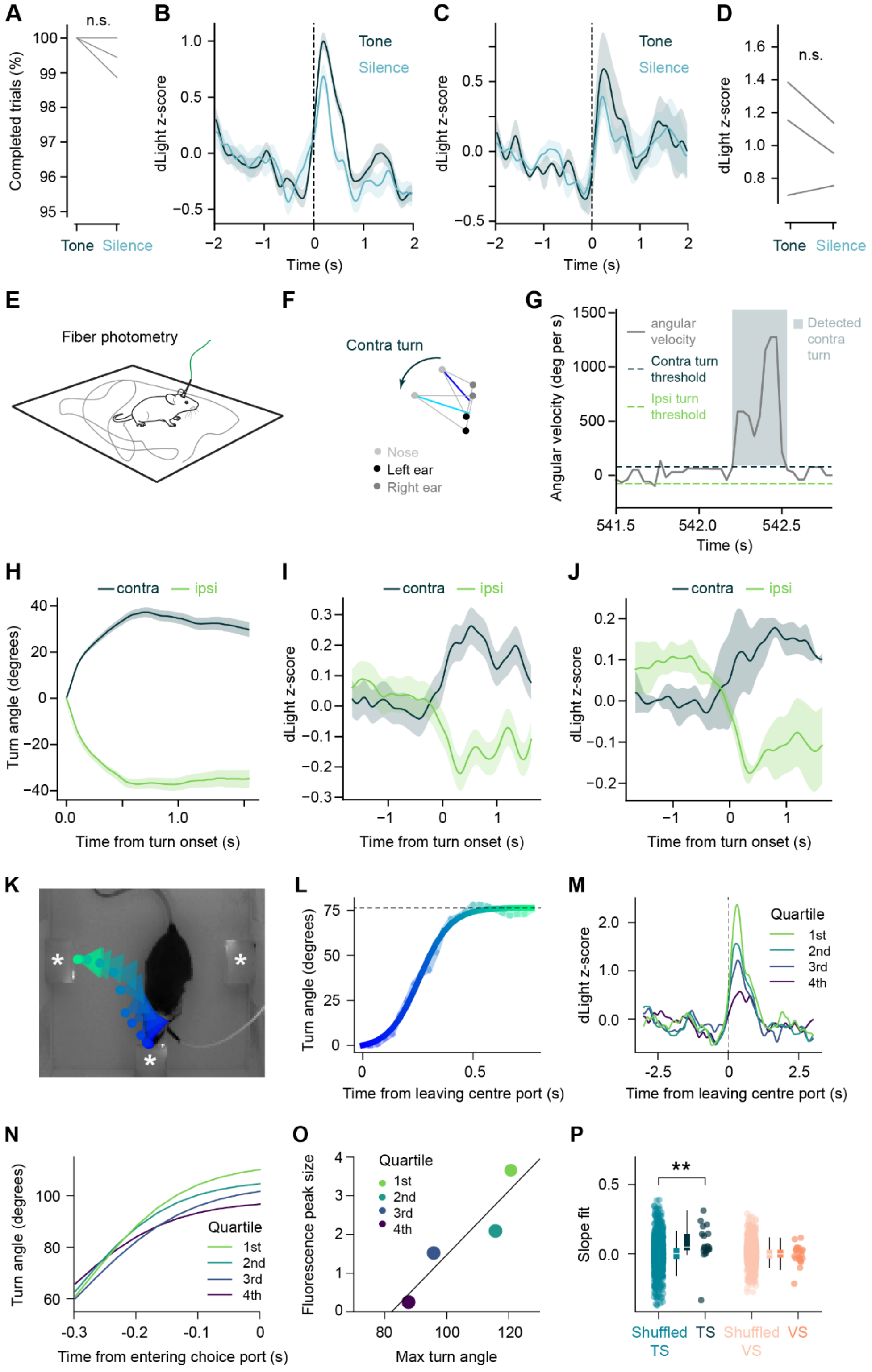


***Figure S4: TS dopamine is related to movement, not cue. A*** *- Change in percentage of completed trials (where animals made a left or right choice after leaving the center port) in trials with a normal stimulus (tone) or silence trials (p=0.23, two related samples t-test).* ***B*** *- dLight responses in the TS to the contralateral choice if it was preceded by a cloud of tones or silence in an example mouse. Traces are aligned to time leaving the center port.* ***C*** *- As for A but average across mice (n=3 mice).* ***D*** *- Change in peak response to the contralateral choice if the associated cloud of tones is replaced with silence (p=0.31, two related samples t-test).* ***E*** *- Animals were allowed to move freely in a different arena to the training box whilst dLight signals were recorded from the TS.* ***F*** *- Head angle during a detected turn in the freely-moving arena. Dark blue line represents the orientation of the head at the*

*beginning of the turn. Light blue line shows head orientation 0.5 s later.* ***G*** *- Contralateral and ipsilateral turns were detected using the angular velocity of the mouse's head. Turns starts were determined based on the angular velocity first exceeding plus or minus 0.5 standard deviations (contra and ipsi).* ***H*** *- Average head angle of an example animal on ipsi and contralateral turns.* ***I*** *- Example dLight response in the TS to contralateral and ipsilateral turn onsets outside of any task in the freely moving area.* ***J -*** *As in I but averaged across animals (n=3 mice).* ***K -*** *Downsampled head position of an animal making a choice on a single trial. The head is represented with a triangle with a circle at the nose.* ***L*** *- Turn angles were calculated using the cumulative angular velocity, with 0 degrees defined as the angle when the animal leaves the center port. Turn angle quantified on a single trial. A sigmoid function was fitted to the turn angle.* ***M*** *- Example traces from a TS recording session separated by size of response in the TS.* ***N*** *- Average turn angle for these quantiles plotted against quantile midpoint (example session).* ***O*** *- The plateau of the sigmoid for each trial turn angle vs the average peak size of the TS photometry signal per quartile based on the photometry signal (example session).* ***P*** *- Data from early in training (first three sessions) is analyzed as shown in E and a regression slope fitted (TS: n = 18 (6 mice, 3 sessions), VS: n = 21 (7 mice, 3 sessions)). The fit slope is then compared to the fit slope if the x and y labels are shuffled in G). The data was shuffled 100 times and the distribution of the fit slopes from the actual data was compared to the shuffled distribution. The same analysis was applied to the VS. For the TS the shuffled and non-shuffled data are significantly different (p=0.001, two-sample t-test). The slopes of the shuffled VS quantiles were not significantly different from the actual data (p=0.55, two-sample t-test).*

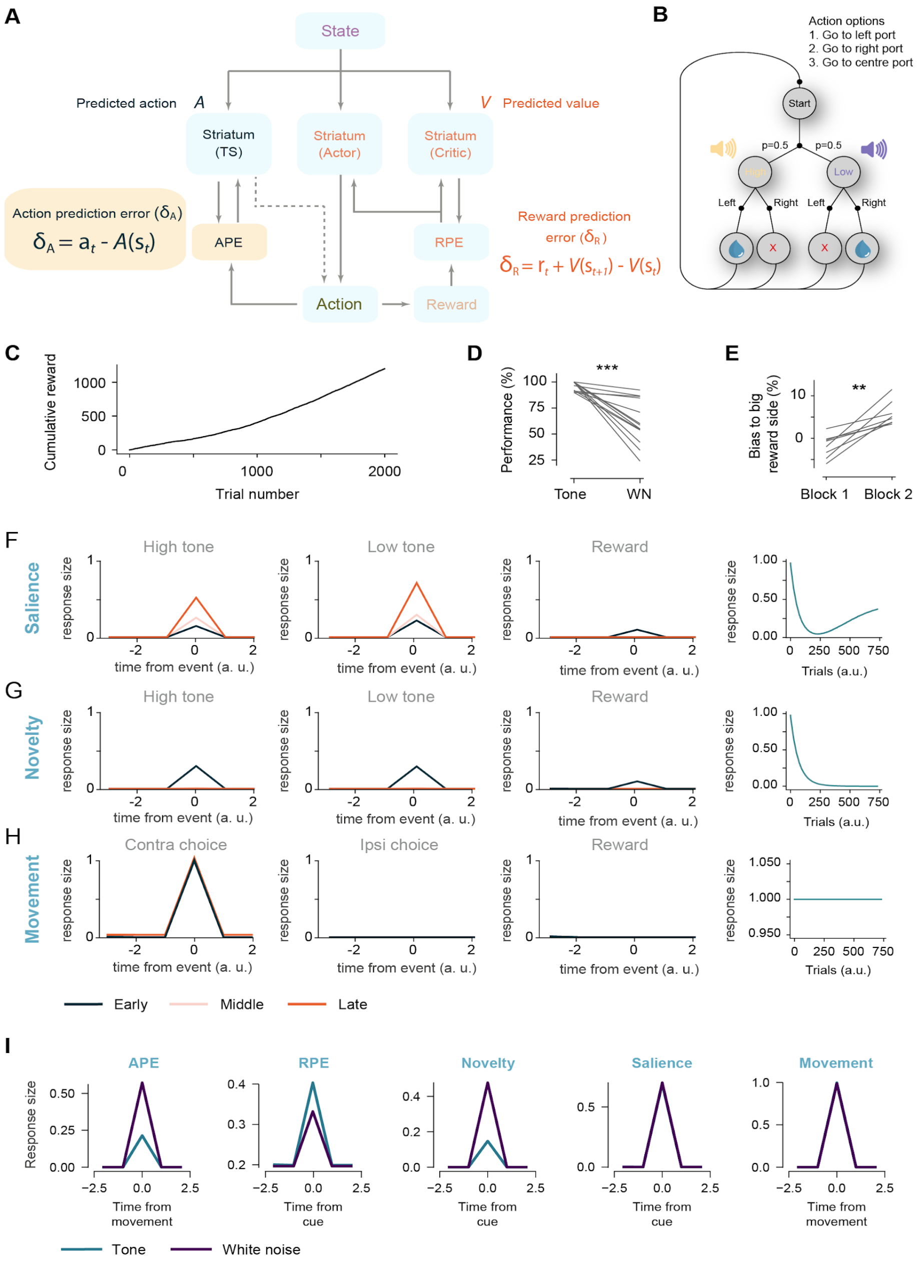


***Figure S5: APE, RPE and other models of dopamine. A** - The model comprises an actor that learns stimulus-action values and guides action choices, a critic that learns a value function that is used to calculate RPE. The RPE signal is broadcast to the actor and critic to update their respective value functions. A value-free system learns to predict actions from those taken in the past and updates its prediction using the difference between its prediction and the action taken (APE). APE and RPE equations are written with respect to time (t), as is common, for illustrative purposes. For the model equations we use dwell time in the state (k) to approximate temporal discounting, see methods. **B** - The Markov decision process used to model the task. **C** - Cumulative rewards received through training. **D** - Performance in the 50 trials before and after the state change (n=13 mice, $p=1.98\times10^{-4}$ two related samples t-test). **E** - Animals' bias to the large reward side before and after the reward is increased in the predicted value change experiment (n=7 mice, p=0.001 two related samples t-test). **F** - Salience model dopamine signal aligned to time of high cue, low cue, and reward. Size of response to cues shown for 100 agents over training. **G** - Novelty model dopamine signal aligned to time of high cue, low cue, and reward. Size of response to cues shown for 100 agents over training. **H** - Movement model dopamine aligned to time of contralateral choice, ipsilateral choice, and reward. Size of response to contralateral choice shown for 100 agents over training. **I** - Model predictions for how the different models of dopamine respond in the state change experiment.*

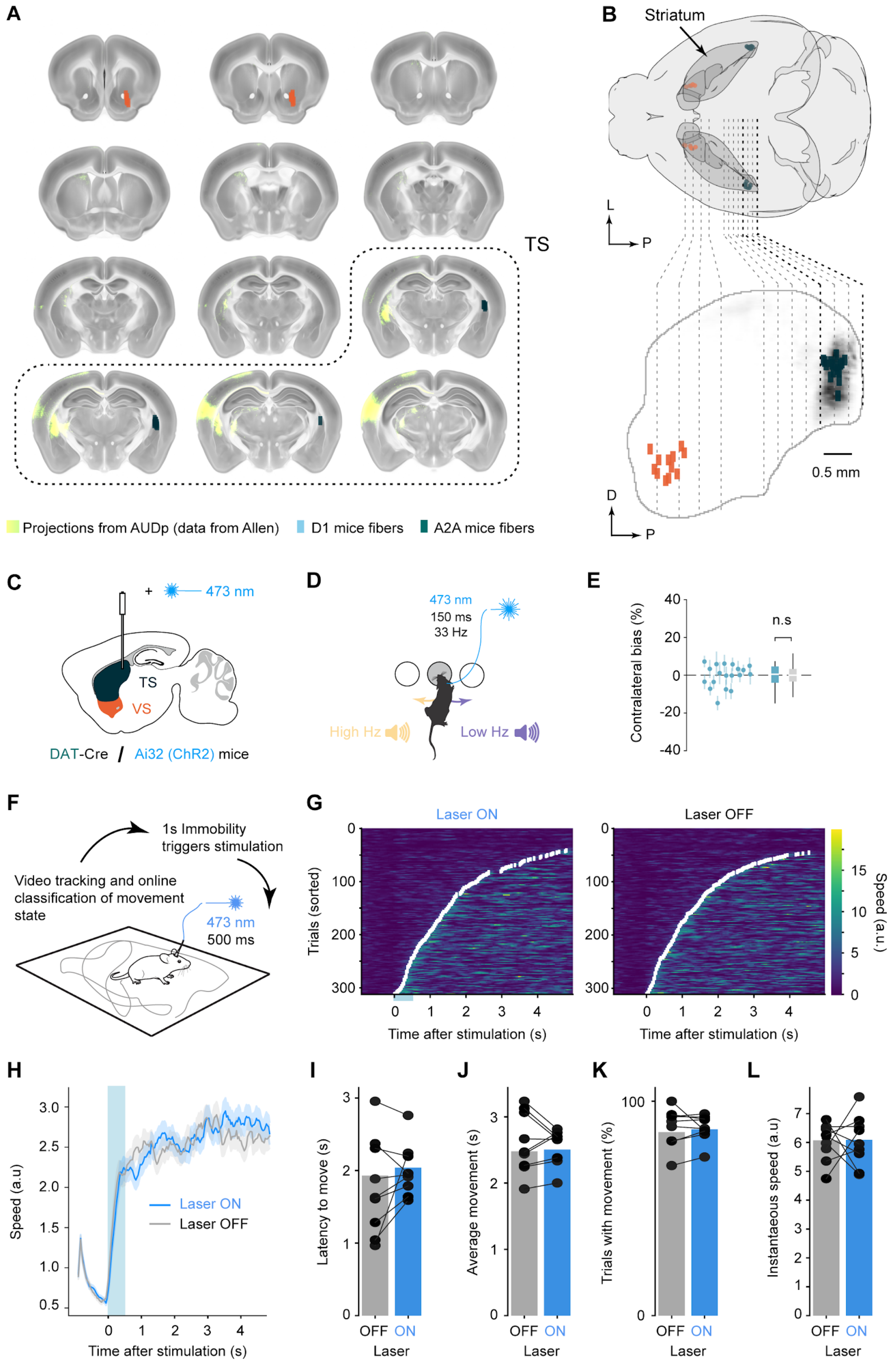
A
TS
Projections from AUDp (data from Allen)
D1 mice fibers
A2A mice fibers
B
Striatum
L
P
D
P
0.5 mm
C
+
473 nm
TS
VS
DAT-Cre / Ai32 (ChR2) mice
D
473 nm
150 ms
33 Hz
High Hz
Low Hz
E
Contralateral bias (%)
n.s
40
20
0
-20
-40
F
1s Immobility triggers stimulation
Video tracking and online classification of movement state
473 nm
500 ms
G
Laser ON
Laser OFF
Trials (sorted)
Time after stimulation (s)
Speed (a.u.)
H
Speed (a.u)
Laser ON
Laser OFF
Time after stimulation (s)
I
Latency to move (s)
OFF ON
Laser
J
Average movement (s)
OFF ON
Laser
K
Trials with movement (%)
OFF ON
Laser
L
Instantaneous speed (a.u)
OFF ON
Laser

***Figure S6: Histology and motor effects of dopamine optostimulation. A*** *- Coronal sections along the striatum indicating fiber placement positions (center tip of the fiber). Note that fibers were inserted in both hemispheres and are mirrored here for illustration purposes. Primary auditory cortex projections are shown in the other hemisphere.* ***B*** *- 3D rendering (top) and side view (bottom) of the same histological data. On the side view, the striatum is outlined and the AUDp projections are indicated in greyscale.* ***C*** *- Schematic of fiber location.* ***D*** *- Schematic of the experiment for E.* ***E*** *- Quantification of the bias (see methods) for each session. Scatter dots represent the means and standard deviation for each session, and coloured box plot represents the distribution of the means compared to the baseline in gray (see methods; p=0.67, Kruskal-Wallis test).* ***F*** *- Schematic of the experiment for G-L.* ***G*** *- Speed heatmap of trials sorted by movement onset for stimulated and control trials (n=9 animals).* ***H*** *- Speed histogram, same data as in G.* ***I, J, K, L*** *- Distribution, as means per mouse, of different movement parameters (see methods) (I: p-value=0.21; J: p-value=0.67; K: p-value=0.87; L: p-value=0.93; t-test on two related samples).*

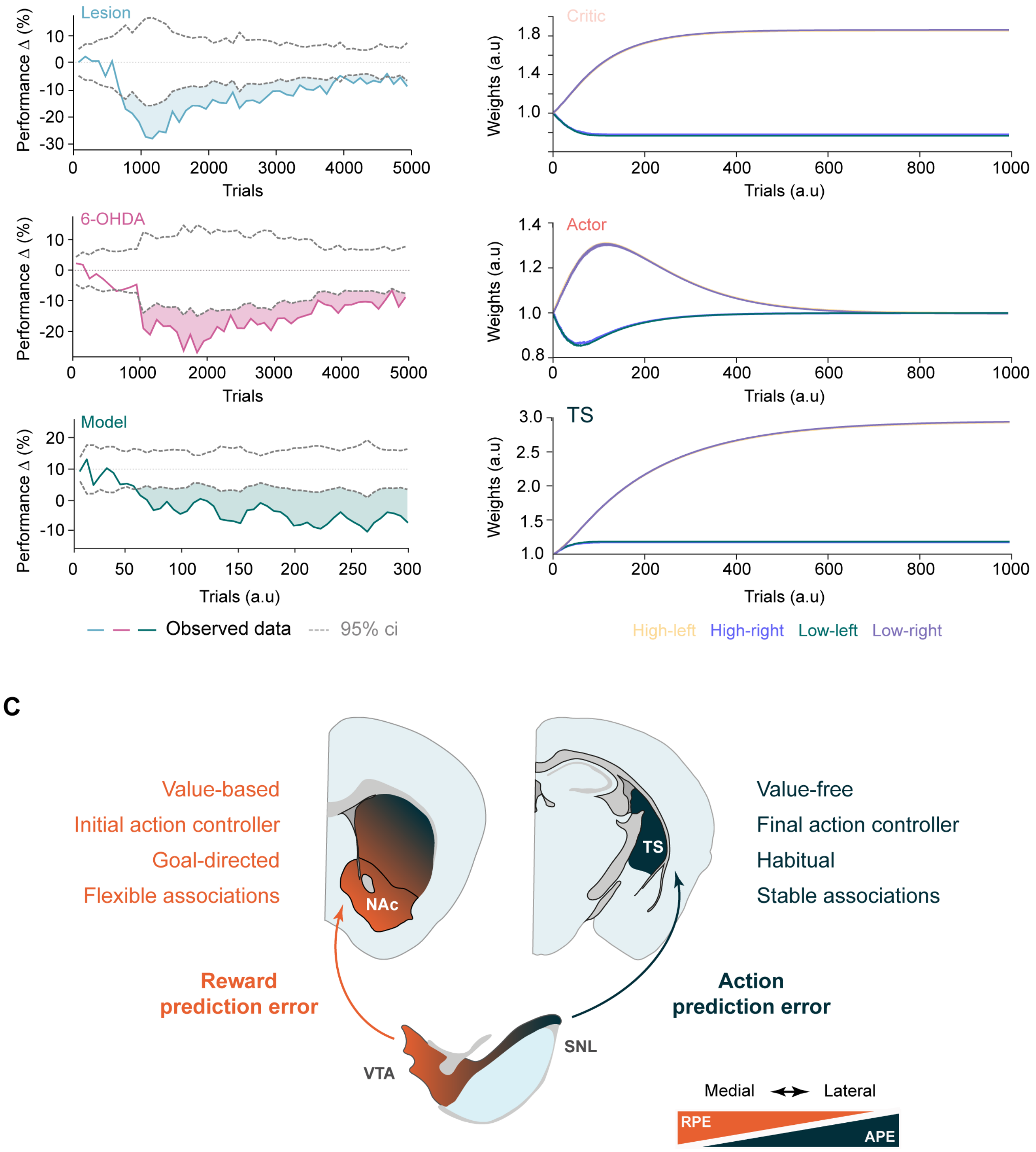


***Figure S7: Network model parameters and summary. A*** *- Moment in learning in which differences in between groups become significant, for the behavioral data and the network model.* ***B*** *- Change of the model weights during learning, as means for 100 agents, for the Critic, Actor, and TS networks.* ***C*** *- Summary schematic of the anatomical representation for the dual-controller model.*
